# Mitochondria serve as a Holdout Compartment for Aggregation-Prone Proteins hindering Efficient Ubiquitin-Dependent Degradation

**DOI:** 10.1101/2025.02.27.640505

**Authors:** Maria E. Gierisch, Enrica Barchi, Mirco Marogna, Moritz H. Wallnöfer, Maria Ankarcrona, Luana Naia, Florian A. Salomons, Nico P. Dantuma

**Affiliations:** Department of Cell and Molecular Biology (CMB), Karolinska Institutet, Solnavägen 9, S-17165 Stockholm, Sweden; Department of Neurobiology, Care Sciences and Society, Division of Neurogeriatrics, Center for Alzheimer Research, Karolinska Institutet, Stockholm, Sweden

**Keywords:** ubiquitin-proteasome system, protein quality control, misfolded proteins, neurodegeneration

## Abstract

The accumulation of protein aggregates has been causatively linked to the pathogenesis of neurodegenerative diseases. In this study, we have conducted a genome-wide CRISPR-Cas9 screen to identify cellular factors that stimulate the degradation of an aggregation-prone reporter protein. Our findings revealed that genes encoding proteins involved in mitochondrial homeostasis, including the translation factor eIF5A, were highly enriched among suppressors of degradation of an aggregation-prone reporter. Conversely, endoplasmic reticulum (ER)-associated ubiquitin ligases facilitated degradation, indicating opposing roles for these cellular compartments in the clearance of aggregation-prone proteins. Genetic or chemical inhibition of eIF5A led to the dissociation of the aggregation-prone substrate from mitochondria, which was accompanied by enhanced degradation through ER-associated ubiquitination. The presence of an aggregation-prone, amphipathic helix that localized the reporter to mitochondria was crucial for the stimulatory effect of eIF5A inhibition. Additionally, the steady-state levels of α-synuclein, a disease-associated protein containing an amphipathic helix that mislocalizes to mitochondria, were reduced upon eIF5A inhibition. We propose that mitochondria behave as a holdout compartment for aggregation-prone proteins, keeping them out of reach of ubiquitin ligases that target them for proteasomal degradation. Therefore, preventing mitochondrial localization of aggregation-prone proteins may offer a viable therapeutic strategy for reducing their levels in neurodegenerative disorders.

## Introduction

Cells are equipped with intricate protein quality control systems that prevent contamination of the intracellular environment by misfolded or otherwise dysfunctional proteins (Sala et al, 2017). These systems capture aberrant proteins and either refold them to their native state or target them for degradation. Given that misfolded proteins are inherently toxic due to their propensity to aggregate, their prompt elimination is critical for cell viability (Bucciantini et al, 2002). Several idiopathic and familial neurodegenerative diseases are linked to the age-dependent accumulation of aggregation-prone proteins in neurons, indicating that inefficient clearance of these aberrant proteins lies at the core of cellular pathology (Soto, 2003). Reducing the steady-state levels of the disease-associated proteins to prevent their accumulation, aggregation and toxicity, presents a promising therapeutic approach. While various strategies have been explored to reduce the synthesis of disease-associated proteins (Alves et al, 2008; Kuijper et al, 2024; Pappada et al, 2022; Tabrizi et al, 2019), leveraging the cell’s intrinsic defense mechanisms against misfolded proteins remains an underexplored avenue.

The ubiquitin-proteasome system (UPS) is primarily responsible for degrading misfolded proteins in the cytoplasmic and nuclear compartments. The two key players are ubiquitin, a protein modifier that tags proteins for degradation (Dikic & Schulman, 2023), and the proteasome, a multi-subunit complex that degrades ubiquitinated proteins (Bhattacharyya et al, 2014). While inefficient clearance of misfolded proteins by the UPS has been proposed as an explanation for the accumulation of ubiquitin-positive protein aggregates in neurodegenerative disorders, recent insights indicate that UPS dysfunction is not necessarily a common feature in the etiology of these diseases (Dantuma & Bott, 2014). For instance, in mouse models for Huntington’s disease (HD) and spinocerebellar ataxia (SCA) type 7, pathogenic aggregation-prone proteins accumulate despite a largely functional UPS (Bett et al, 2009; Bowman et al, 2005; Maynard et al, 2009). The presence of an operational UPS in neurons that succumb to toxic protein aggregates raises the question of whether this protective mechanism can be therapeutically harnessed to more effectively target disease-associated proteins.

The UPS has gained interest as a potential therapeutic target due to the numerous druggable proteins involved in this proteolytic pathway. Most efforts have focused on developing drugs that inhibit the UPS, some of which are now clinically used for treating of hematological malignancies (Goldberg, 2012). However, for neurodegenerative disorders, stimulating UPS activity to degrade aggregation-prone proteins presents a more challenging goal, as drugs typically exert their therapeutic effects through inhibition rather than stimulation of enzymatic activities. Cells possess complex regulatory circuits that can enhance UPS capacity in response to acute increases in demand, such as during proteotoxic stress (Sala et al, 2017). This adaptability suggests that the UPS, when unchallenged, may not operate at its maximal capacity, indicating potential for pharmaceutical stimulation. A few targets for stimulating UPS activity have been identified (Lee et al, 2010; Leestemaker et al, 2017; Matilainen et al, 2013), providing proof-of-principle that UPS activity can be enhanced. However, given the enormous complexity of this proteolytic system and the large number of proteins involved, the main challenge lies in identifying viable drug targets.

Mislocalization of aggregation-prone proteins to mitochondria has been reported for several proteins associated with neurodegenerative diseases. For example, α-synuclein (Devi et al, 2008), mutant huntingtin (Orr et al, 2008), and amyloid β-peptide (Hansson Petersen et al, 2008) have been found to localize at the mitochondrial compartment in cellular and animal models for neurodegeneration. The tendency of these proteins to bind mitochondria may be intrinsically linked to their hydrophobic nature, which also contributes to their propensity to form insoluble aggregates. While many studies have investigated the role of mislocalized proteins in mitochondrial dysfunction, little is known about the effect of mitochondrial sequestration of aggregation-prone proteins on cell’s ability to clear these proteins through intracellular protein degradation. Efficient proteasomal degradation depends on the proximity of the aggregation-prone proteins to the ubiquitin ligases that target them for degradation. Therefore, the subcellular localization of these proteins may affect their degradation kinetics and modulate their accumulation in affected cells.

Here, we report on a genome-wide screen to identify cellular factors that regulate the ubiquitin-dependent proteasomal degradation of aggregation-prone proteins. Our findings indicate that biochemical or genetic inhibition of the translation elongation factor eIF5A, a regulator of mitochondrial homeostasis, accelerates the degradation of aggregation-prone proteins that contain amphipathic helices and mislocalize to mitochondria. Our data suggest that the sequestration of aggregation-prone proteins to mitochondria impedes degradation, as this largely depends on ubiquitinating enzymes localized at the endoplasmic reticulum (ER). We propose that preventing the sequestration of disease-associated proteins in the protective mitochondrial environment can stimulate their degradation and identified eIF5A inhibition as a potential target for this purpose.

## Results

### Generation of reporter cell line for the CRISPR-Cas9 screen

Designing a CRISPR-Cas9 screen to identify regulators of proteasomal degradation of aggregation-prone proteins using a reporter substrate involves several technical challenges. First, the inherent toxicity of protein aggregates complicates the stable expression of aggregation-prone proteins necessary for a phenotypic screen (Bucciantini et al, 2002). Second, the expression levels of these proteins must allow for detection of both an increase and decrease in the steady-state levels. Third, changes in steady-state levels should result from enhanced degradation and be distinguishable from changes due to reduced synthesis. To address these issues, we utilized the commonly used aggregation-prone reporter yellow fluorescent protein (YFP)-CL1, which can be stably expressed in cells without overt signs of toxicity (Bence et al, 2001). Under physiological conditions, YFP-CL1 is distributed throughout the cytosol and nucleus at low but detectable levels, but it accumulates and forms intracellular aggregates when degradation is impaired (Menendez-Benito et al, 2005; Salomons et al, 2009). Aggregation is mediated by the CL1 degradation signal, which consists of an aggregation-prone amphipathic helix (Gilon et al, 2000). Flow cytometric analysis showed that the unrelated proteasome inhibitors epoxomicin and bortezomib as well as the E1 ubiquitin activase inhibitor TAK-243 induced a very similar accumulation of YFP-CL1 in a human melanoma MelJuSo cell line (**Fig. 1A)**. Consistent with the exclusive role of the UPS in degrading YFP-CL1, administration of the autophagy inhibitor Bafilomycin A1 did not affect the steady-state levels of the reporter (**Fig. 1A)**.

**Figure 1:**
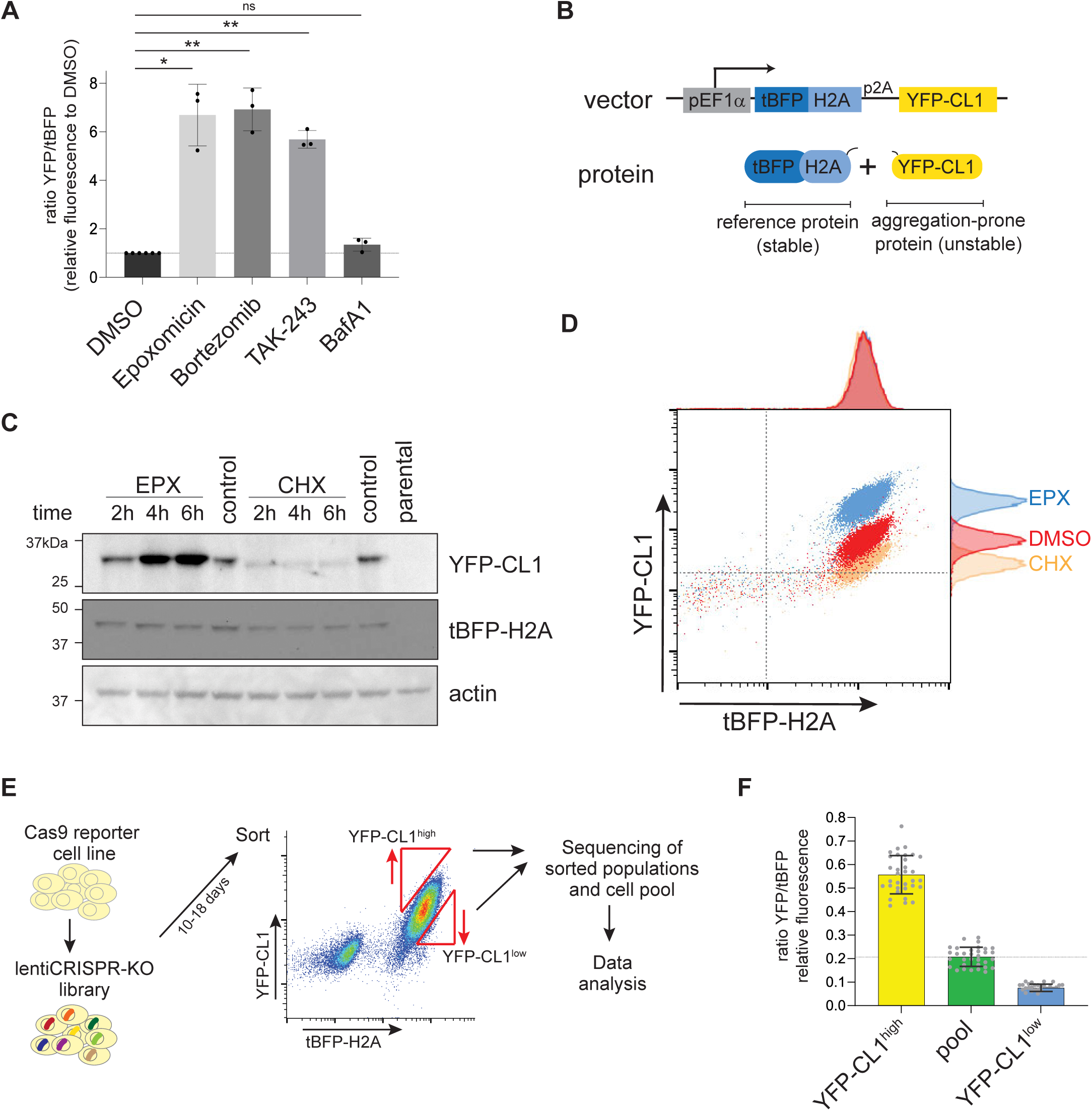
Generation of reporter cell line for the CRISPR-Cas9 screen. **(A)** MelJuSo YFP-CL1 reporter cells were plated for 72 hours and treated with the proteasome inhibitors epoxomicin (100 nM) and bortezomib (25 nM), the E1 inhibitor TAK-243 (10 µM) or the autophagy inhibitor BafA1 (100 nM) for the last 8 hours and analyzed by flow cytometry (n=3, mean ± SD, one-sample t-test, *P<0.05, **P<0.01, ns: non-significant). **(B)** Schematics of the dual-fluorescent reporter used for the screening. **(C)** MelJuSo cells stably expressing the dual-fluorescent reporter were treated with 100 nM epoxomicin (EPX) or 20 µg/ml cycloheximide (CHX) for the indicated timings and analyzed by western blotting. **(D)** Stable dual-fluorescent reporter cells were plated for 24 hours and treated with 100 nM epoxomicin (EPX) or 20 µg/ml cycloheximide (CHX) for the last 7 hours and analyzed by flow cytometry. **(E)** Screening outline, Cas9 dual-fluorescent reporter cells were transduced with a lentiviral CRISPR library at an MOI∼0.4 and cultured. The 0.5% populations with the highest (YFP-CL1^high^) and lowest (YFP-CL1^low^) YFP-CL1/tBFP-H2A ratios were sorted and subjected for sequencing. **(F)** Post-sort analysis of the sorted cells when gating for the requirements from (E). Dots represent the YFP/tBFP ratio at the start of each 2-hour long sort, mean ± SD.

To distinguish in our screen between changes in protein levels due to degradation versus protein synthesis, we used the stable reference protein histone H2A tagged with tag blue fluorescent protein (tBFP), which was expressed from the same transcript using a p2A ribosome-skipping motif, allowing quantification of the reporter as the ratio between YFP and tBFP fluorescence (**Fig. 1B**). The reporter construct was used to generate a MelJuSo cell line stably expressing tBFP-H2A-p2A-YFP-CL1 from an EF1α promoter. Incubation with the translation inhibitor cycloheximide confirmed that the YFP-CL1 was rapidly degraded, whereas the tBFP-H2A was long-lived (**Fig. 1C**). Moreover, treatment with the proteasome inhibitor epoxomicin increased the levels of YFP-CL1 but not of the long-lived tBFP-H2A. Microscopic analysis of cells treated with epoxomicin or TAK-243 showed increased YFP-CL1 fluorescence and the formation of perinuclear foci, confirming its aggregation-prone nature (**Suppl. Fig. 1A, B**). Flow cytometric analysis of the stable cell line showed that reduced or elevated levels of YFP-CL1 upon cycloheximide and epoxomicin treatment, respectively, could be quantified relative to the tBFP-H2A reference protein, allowing for sorting for high and low expressing populations based on YFP/tBFP ratio (**Fig. 1D**).

We then designed a pipeline for a fluorescence-activated cell sorting (FACS)-based genome-wide CRISPR-Cas9 screen to identify positive and negative modulators of YFP-CL1 degradation (**Fig. 1E**). Cas9-expressing reporter cells were transduced with a lentiviral library consisting of the Brunello knockout sgRNA library that included Unique Molecular Identifiers (UMIs) (Doench et al, 2016). This pooled library has genome-wide coverage and consists of four sgRNAs targeting each gene in the human genome. After culture for 10-18 days, approximately 0.5% of the cells exhibiting the highest and lowest YFP/tBFP ratios were sorted by FACS, resulting in the YFP-CL1^high^ and YFP-CL1^low^ populations, respectively (**Fig. 1F**). The enriched sgRNAs in these populations were identified by next generation sequencing of the UMIs.

### Identification of proteins stimulating in YFP-CL1 degradation

We first analyzed the YFP-CL1^high^ population to validate our screening procedure using the well-characterized degradation pathway of this reporter (Gilon et al, 1998; Gilon et al, 2000; Leto et al, 2019; Stefanovic-Barrett et al, 2018). Analysis of the YFP-CL1^high^ population using the Model-based Analysis of Genome-wide CRISPR/Cas9 Knockout (MAGeCK) method (Li et al, 2014) and by UMI lineage dropout method (Schmierer et al, 2017) showed that for 1574 genes at least two sgRNAs were significantly enriched over the whole population (**Suppl. Table 1**). Of these sgRNAs-targeting 824 genes were enriched in both replicates (**Fig. 2A**). Gene ontology (GO) annotation showed that the largest GO classes consisted of genes linked to the insertion of membrane proteins into the endoplasmic reticulum (ER) as well as protein ubiquitination and degradation (**Fig. 2B**). This is consistent with earlier studies showing that the engineered CL1 degradation signal is an aggregation-prone amphipathic helix resembling the membrane targeting sequences of tail-anchored proteins (Stefanovic-Barrett et al, 2018) and primarily engages ER-associated ubiquitination enzymes (Gilon et al, 1998; Gilon et al, 2000; Leto et al, 2019). Among the enriched genes in the YFP-CL1^high^ population we found the ubiquitin conjugase UBE2G2 together with AUP1, which recruits UBE2G2 to the ER membrane (Spandl et al, 2011), the ubiquitin ligase RNF139 (also known as TRC8), implicated in ER-associated degradation (ERAD) (Krshnan et al, 2022) and the ubiquitin elongation factor UBE3C, which extends ubiquitin chains on YFP-CL1 with K29-linked ubiquitin (Michel et al, 2015), consistent with earlier reports (Leto et al, 2019; Stefanovic-Barrett et al, 2018) (**Fig. 2C**). Depletion of these ubiquitination enzymes confirmed their role in regulating YFP-CL1 stability (**Fig. 2D, Suppl. Fig. 2A-C**). The importance of ER localization for efficient proteasomal degradation of YFP-CL1 was further supported by the enrichment of sgRNAs targeting EMC1, EMC2, EMC6, EMC8, and MMGT1, which are part of the ER membrane complex (EMC) that facilitates membrane insertion of tail-anchored proteins (Zhu et al, 2024). Furthermore, the enriched sgRNA-targeted genes also included the ER-associated protein SRP1, which targets proteins to the ER (Aviram & Schuldiner, 2017).

**Figure 2:**
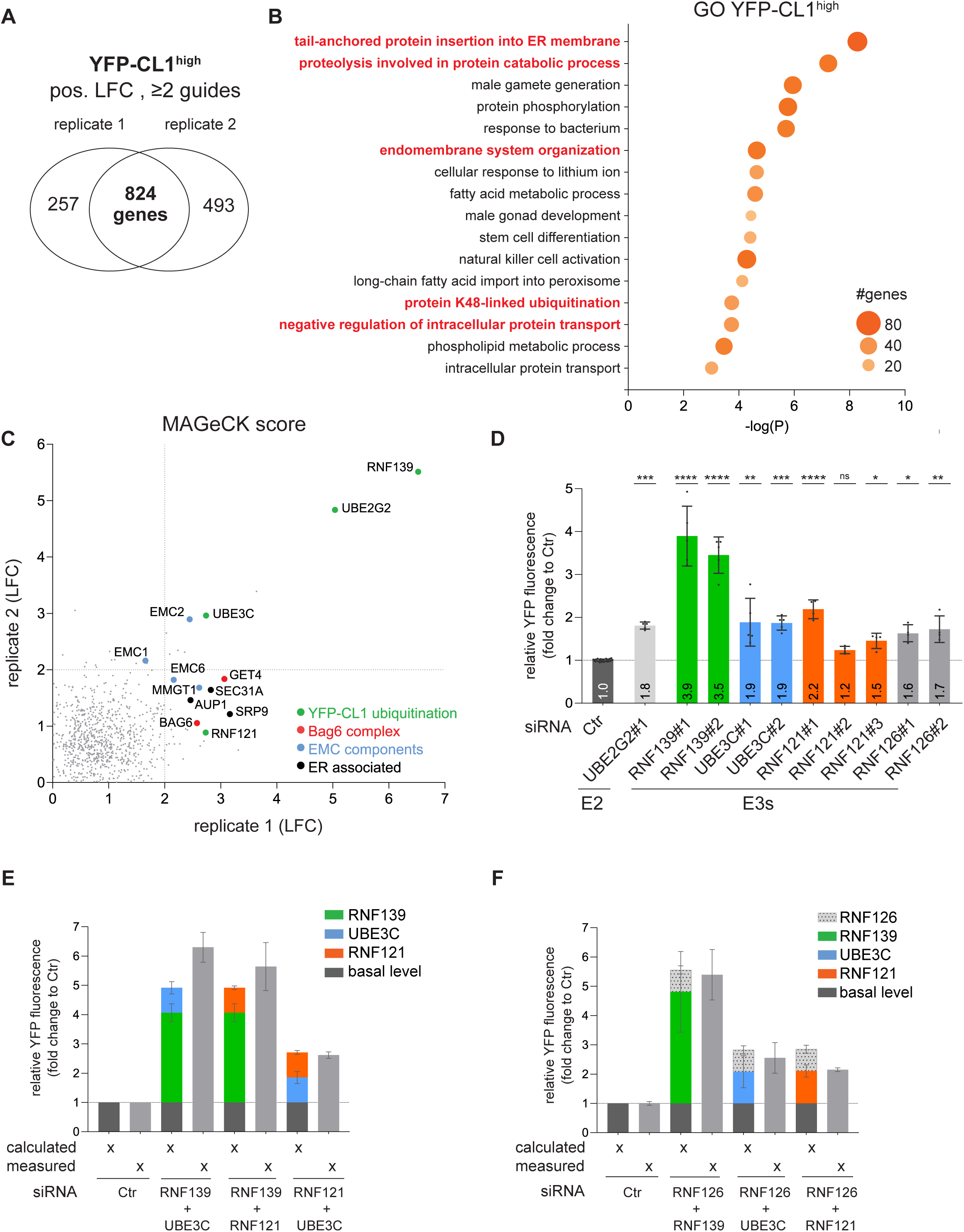
Identification of proteins stimulating in YFP-CL1 degradation. **(A)** SgRNA enrichment of the YFP-CL1^high^ population over the pool in the two independent replicates with the indicated criteria. **(B)** Gene Ontology (GO) analysis of 824 genes that overlap between the two replicates. **(C)** The log fold change (LFC) of the enriched genes of the two replicates were plotted against each other and selected pathways are color-coded. **(D)** MelJuSo YFP-CL1 reporter cells were transiently transfected with siRNAs for 72 hours and analyzed by flow cytometry for YFP fluorescence (n=5 for RNF139 andUBE3C, n=4 forRNF121, RNF126 and UBE2G2), mean ± SD, Kruskal-Wallis test with uncorrected Dunn’s multiple comparisons, *P<0.05, **P<0.01, ***P<0.001, ****P<0.0001, ns: non-significant). (**E, F**) MelJuSo YFP-CL1 reporter cells were transfected with 10 nM siRNA against single E3 ligases together with 10 nM siCtr or with a combination of two E3 ligase siRNAs (each 10 nM) for 72 hours and analyzed by flow cytometry for YFP fluorescence. The calculated bar presents the YFP-CL1 fluorescence of the theoretical combination of the single measured E3 siRNAs while the measured bar presents the actual E3 siRNA combination (n=3, mean ± SD).

An alternative known pathway for YFP-CL1 degradation involves the BAG6-GET1 chaperone complex (Minami et al, 2010), which intercepts soluble tail-anchored proteins and either targets them for membrane (re-)insertion or presents them to RNF126 for ubiquitination and subsequent proteasomal degradation (Hessa et al, 2011; Rodrigo-Brenni et al, 2014). While the sgRNAs-targeting BAG6 and GET1 were enriched, those targeting RNF126 were not found in the YFP-CL1^high^ population. However, siRNA-depletion confirmed that RNF126 promotes YFP-CL1 degradation, albeit less prominent than RNF139 (**Fig. 2D, Suppl. Fig. 2D**).

The identification of several expected hits validated the screen, but new candidates were also discovered. These included RNF121, a ubiquitin ligase not previously implicated in targeting YFP-CL1 or other tail-anchored proteins for degradation. Given its localization at the cytosolic face of the ER membrane and its role in degrading a voltage-gated sodium channel (Ogino et al, 2015), RNF121 fits the profile of a YFP-CL1-targeting ubiquitin ligase. Immunostaining confirmed a distribution of RNF121 resembling that of ER-associated proteins (**Suppl. Fig. 2E**), which was further corroborated by subcellular fractionation where RNF121 was detected in ER fractions (**Supp. Fig. 2F**). Depletion of RNF121 increased YFP-CL1 levels, although not to the same extent as RNF139 depletion (**Fig. 2D, Suppl. Fig. 2G, H**). Co-depletion of RNF139 + UBE3C, RNF139 + RNF121, and RNF121 + UBE3C caused an increase in YFP-CL1 that matched or exceeded the predicted additive effect of single depletions, suggesting that these ubiquitination enzymes may independently target YFP-CL1 for proteasomal degradation (**Fig. 2E, Suppl. Fig. 2I**). While co-depletion of RNF126 and RNF139 or RNF126 and UBE3C suggested independent targeting mechanisms, the combined depletion of RNF126 and RNF121 did not have an additive effect, implying that the ligases may act on different steps of the same pathway (**Fig. 2F, Suppl. Fig. 2J**). This is reminiscent of earlier data indicating that the degradation of some ER-associated substrates requires separate ubiquitination events for ER extraction and proteasomal degradation (Hu et al, 2020).

The presence of several ubiquitin ligases previously implicated in YFP-CL1 degradation within the YFP-CL1^high^ population validates our screen and shows that our experimental setup can effectively identify modulators of YFP-CL1 degradation. Additionally, we identified RNF121 as a novel ER-associated ubiquitin ligase involved in the stability of tail-anchored substrates. The identification of several ER-associated ubiquitin ligases that target YFP-CL1 combined with the absence of ubiquitin ligases from other membranous organelles supports a model where the cytosolic face of the ER is the predominant site for ubiquitination of this aggregation-prone reporter substrate.

### Identification of proteins suppressing YFP-CL1 degradation

We next focused on the sgRNAs enriched in the YFP-CL1^low^ population (**Suppl. Table 1**). Compared to the YFP-CL1^high^ data set, a larger number of enriched genes were found in the two replicates (1187 genes) (**Fig. 3A),** but with an overall lower average enrichment score (**Suppl. Table 2**). The enriched genes included several encoding proteins regulating exit from ER, suggesting that increased ER residence time may correlate with a decreased stability of YFP-CL1 (**Fig. 3B**). The GO annotation indicated a link to the mitochondrial compartment as several of the largest GO classes shared a connection to mitochondrial homeostasis (**Fig. 3B**). Using MitoCarta for more detailed annotation (Rath et al, 2021), we found that enriched sgRNAs were directed against genes involved in mitochondrial RNA metabolism, translation and import of mitochondrial proteins, as well as genes encoding components of the respiratory chain complexes I, IV and V (**Fig. 3C**). Notably, the translation factor eIF5A, a general regulator of mitochondrial homeostasis (Barba-Aliaga & Alepuz, 2022), was the only gene that was strongly enriched in both replicates (**Fig. 3D**).

**Figure 3:**
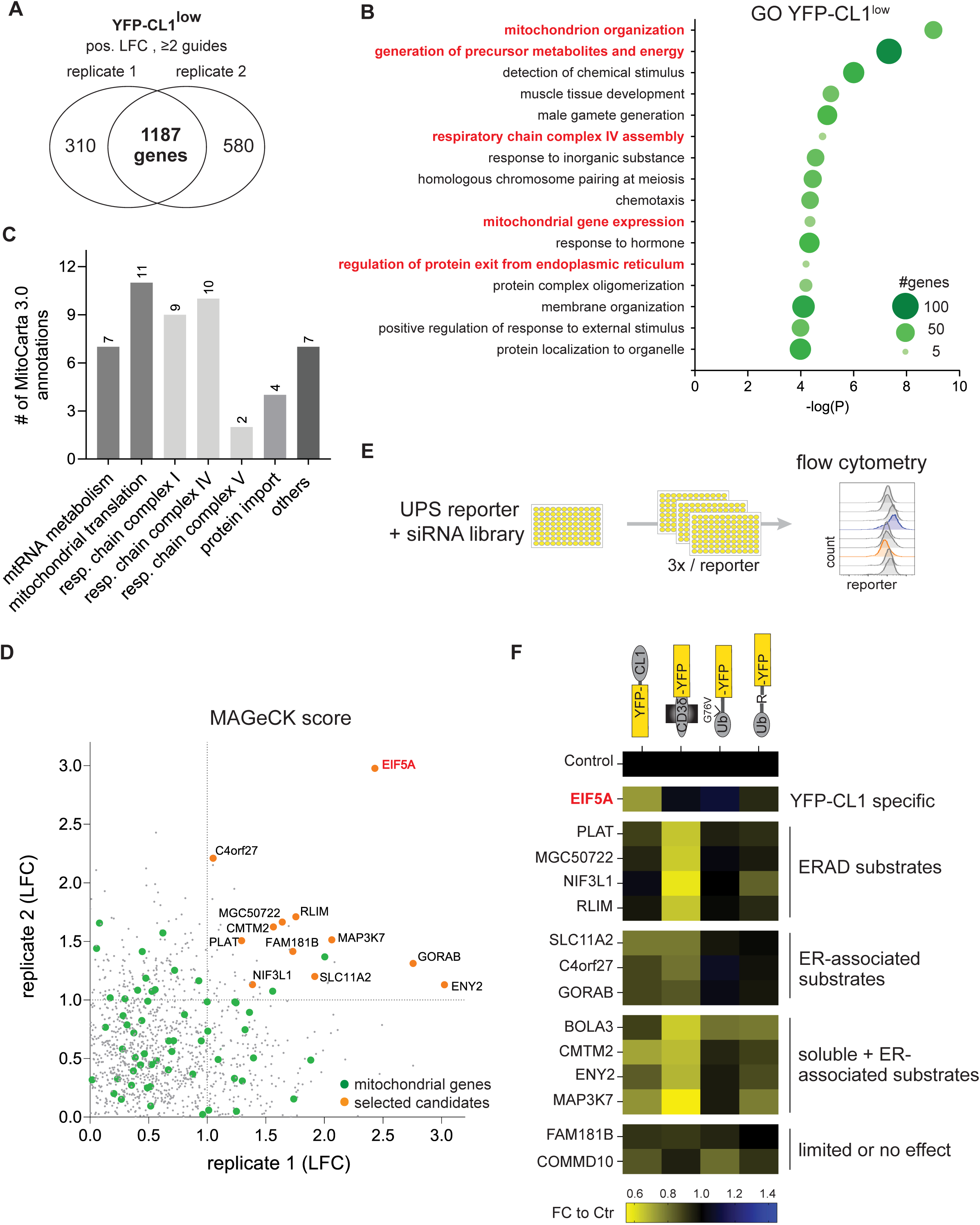
Identification of proteins suppressing of YFP-CL1 degradation. **(A)** SgRNA enrichment of the YFP-CL1^low^ population over the pool in the two independent replicates with the indicated criteria. **(B)** Gene Ontology (GO) analysis of 1187 genes that overlap between the two replicates. **(C)** MitoCarta 3.0 was used to map the enriched mitochondrial genes to distinct groups. **(D)** The log fold change (LFC) of the enriched genes of the two replicates were plotted and mitochondrial genes are indicated in green. Top enriched candidates for further evaluation are indicated orange. **(E)** Outline of the siRNA-based mini-screen. **(F)** MelJuSo reporter cells were transiently transfected with siRNAs against the selected candidates for 72 hours and analyzed by flow cytometry for YFP fluorescence. Each square presents pooled data of three independent screening rounds with three different siRNAs calculated as a fold change to the control. Genes are clustered into five different groups.

We then performed a screen with siRNA targeting eIF5A and nine additional candidates scoring the highest positive logarithm fold change (LFC) based on the MAGeCK analysis and were identified by at least two sgRNAs in both replicates. Two more genes that ranked high in the second replicate of the MAGeCK analysis (BOLA3 and COMMD10) were also included. We determined the effect of depletion of these thirteen candidates on the steady-state levels of four different classes of UPS reporter substrates including, in addition to YFP-CL1, the ERAD substrate CD3δ-YFP, the soluble ubiquitin-fusion degradation (UFD) substrate Ub^G76V^-YFP and the soluble N-end rule substrate Ub-R-YFP (Menendez-Benito et al, 2005). Due to the transient nature of siRNA depletion, reporter levels were already determined after 72 hours, which is considerably shorter than the 10-18 days after sgRNA transduction employed in the genetic screen (**Fig. 3E**). Despite the shorter time span, a >25% reduction in the levels of the YFP reporters was observed upon the depletion of several candidates. Interestingly, only the depletion of eIF5A selectively reduced the steady-state levels of YFP-CL1 without a similar substantial decrease in the levels of ERAD and soluble substrates (**Fig. 3F**).

To our surprise, none of the candidates previously identified as suppressors of the UPS were found to be enriched in the YFP-CL1^low^ population. To determine whether this was due to the non-saturating conditions of the screen or to the experimental model used, we analyzed the impact of depleting several known suppressors on the degradation of the YFP-CL1 reporter. Notably, siRNA depletion of USP14 (Lee et al, 2010), UCH-L5 (Matilainen et al, 2013), and p38 (Leestemaker et al, 2017), which have been found to stimulate proteasomal degradation in other experimental models, did not cause a reduction of YFP-CL1 (**Suppl. Fig. 3A-C**). This suggests that their effects may be substrate- and/or cell-type specific. Therefore, we conclude that among our candidates, only depletion of the elongation factor eIF5A selectively reduces the steady-state levels of the YFP-CL1 reporter.

### Depletion or inhibition of eIF5A selectively reduces YFP-CL1 levels

Considering the selective effect of eIF5A depletion on YFP-CL1 steady-state levels, we focused our efforts on this candidate. Treatment of cells with three independent siRNAs, which efficiently knocked down eIF5A expression (**Suppl. Fig. 4A**), confirmed that eIF5A depletion significantly reduced YFP-CL1 levels (**Fig. 4A**) without affecting the soluble substrate Ub^G76V^-YFP (**Fig. 4B**).

**Figure 4:**
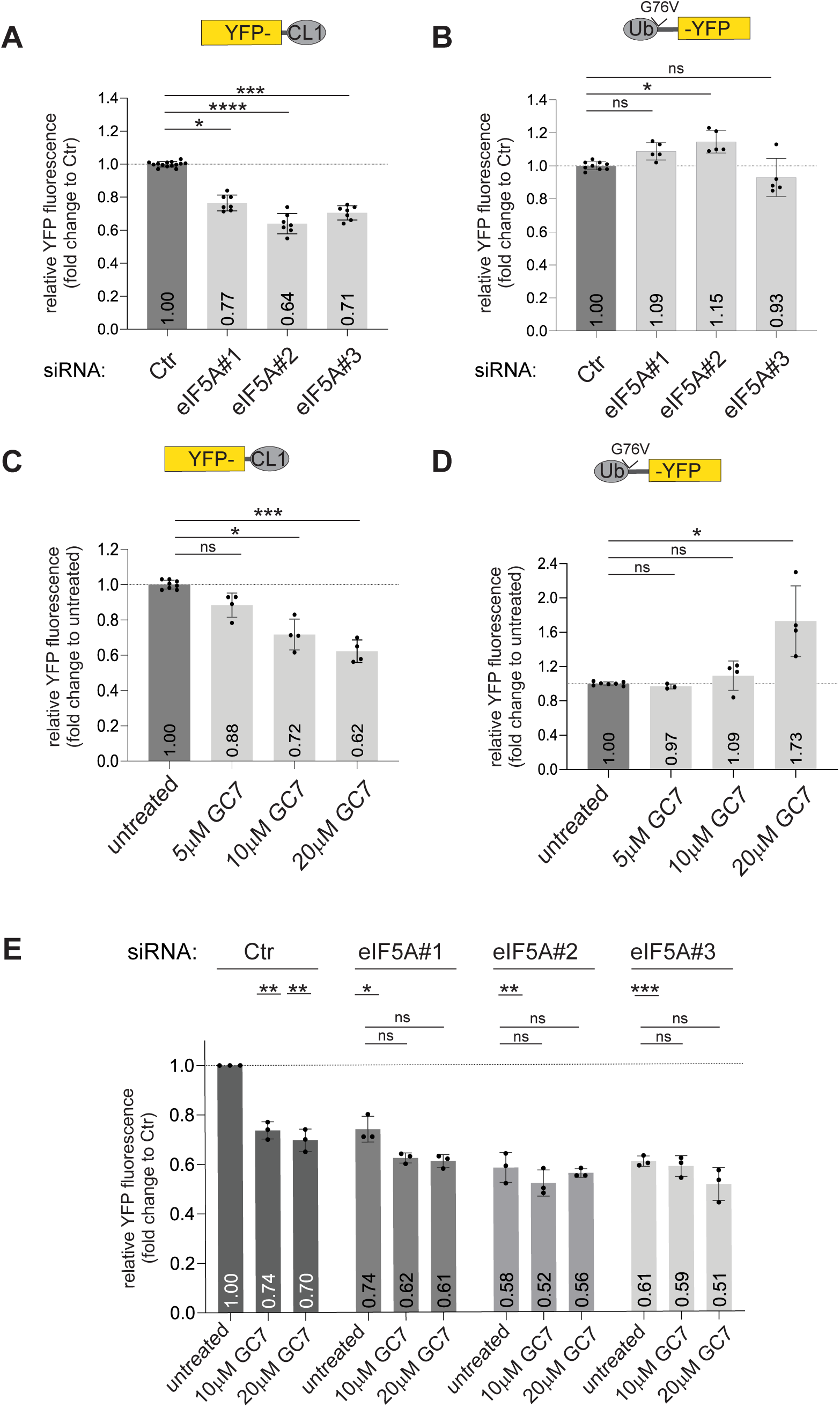
Depletion or inhibition of eIF5A selectively reduces YFP-CL1 levels. **(A, B)** MelJuSo reporter cells YFP-CL1 (A) and Ub^G76V^-YFP (B) were transfected with 20 nM siRNAs against eIF5A for 72 hours and analyzed by flow cytometry for YFP fluorescence (n=7 (A), n=5 (B), mean ± SD, Kruskal-Wallis test, *P<0.05, ***P<0.001, ****P<0.0001, ns: non-significant). **(C, D)** MelJuso reporter cells for YFP-CL1 (C) and Ub^G76V^-YFP (D) were treated with GC7 for the indicated concentrations for 24 hours and analyzed by flow cytometry for YFP fluorescence (n=4, mean ± SD, Kruskal-Wallis test, *P<0.05, ***P<0.001, ns: non-significant). **(E)** MelJuSo YFP-CL1 cells were transfected with 20 nM siRNAs against EIF5A for 72 hours, treated the last 24 hours with the 10 µM or 20 µM GC7 and analyzed by flow cytometry for YFP fluorescence (n=3, mean ± SD, one sample t-test when compared to siCtr or Kruskal-Wallis test when samples were compared within each siEIF5A group, *P<0.05, **P<0.01, ***P<0.001).

For its physiological activity in translation, eIF5A strictly depends on a posttranslational modification that converts a specific lysine residue into a hypusine, a process mediated by the successive action of deoxyhypusine synthase (DHS) and deoxyhypusine hydroxylase (DOHH) (Park et al, 2022). Since eIF5A is the only known hypusination substrate, selective inhibition of eIF5A can be achieved by blocking enzymes involved in hypusination (Jakus et al, 1993). Indeed, treatment with the DHS inhibitor N1-guanyl-1,7-diaminoheptane (GC7) efficiently prevented eIF5A hypusination (**Suppl. Fig. 4B**). Importantly, treatment with GC7 also phenocopied the effect of eIF5A depletion by inducing a dose-dependent reduction in the levels of YFP-CL1 (**Fig. 4C**). In contrast, the levels of Ub^G76V^-YFP were unaffected and even elevated at the highest GC7 concentration (**Fig. 4D**). GC7 administration did not affect the expression levels of YFP-CL1 in eIF5A-depleted cells, confirming that the effect of GC7 on YFP-CL1 levels is executed through inhibition of eIF5A hypusination (**Fig. 4E, Suppl. Fig. 4C**).

### Enhanced ubiquitin-dependent proteasomal degradation is responsible for reduced YFP-CL1 levels in eIF5A-deficient cells

The effect of eIF5A depletion on YFP-CL1 levels was abolished by treatment with the proteasome inhibitors epoxomicin or bortezomib, or by blocking ubiquitination with TAK-243, but was not affected by inhibition of autophagy with Bafilomycin A1 (**Fig. 5A**). This confirms that enhanced degradation, rather than reduced synthesis, is responsible for the reduced YFP-CL1 levels. It also supports the conclusion that eIF5A depletion promotes ubiquitin-dependent proteasomal degradation and argues against a role for autophagy in the enhanced clearance of YFP-CL1.

**Figure 5:**
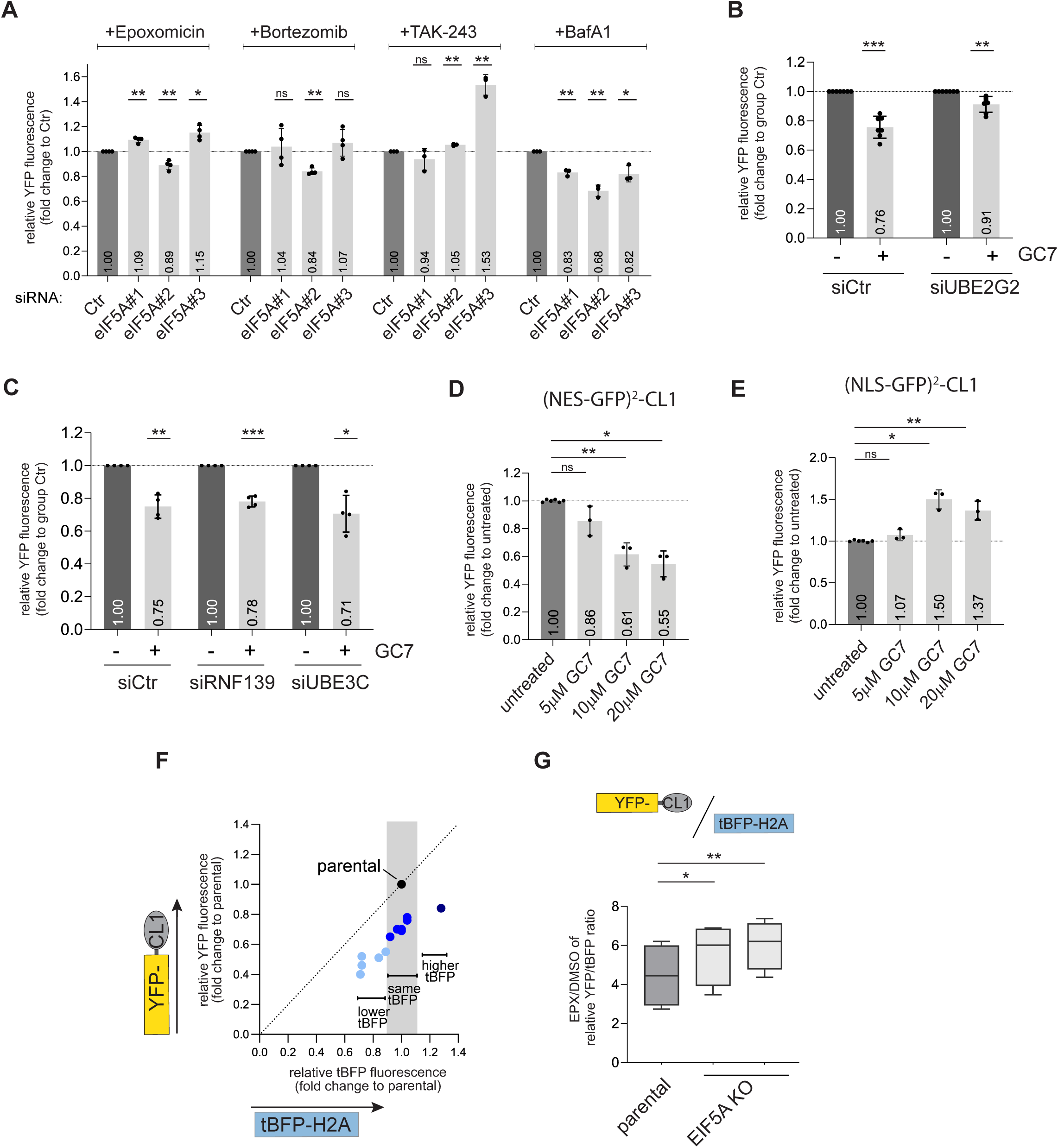
Enhanced ubiquitin-dependent proteasomal degradation is responsible for reduced YFP-CL1 levels in eIF5A-deficient cells>. **(A)** MelJuSo reporter cells expressing YFP-CL1 were transfected with 20 nM siRNAs against eIF5A for 72 hours. The final 8 hours cells were incubated with 100 nM epoxomicin, 25 nM bortezomib, 10 µM TAK-243 or 100 nM BafA1. (n=4 for EPX and bortezomib // n=3 for TAK-243and BafA1, mean ± SD, one-sample t-test, *P<0.05, **P<0.01, ns: non-significant). **(B, C)** MelJuSo stable YFP-CL1 cells were transfected with 20 nM siRNA for 72 hours and treated the last 24 hours with 12 µM GC7. Samples were analyzed by flow cytometry for YFP expression and each siRNA condition was related separately to their respective control (n=7 for (B) and n=4 for (C), mean ± SD, one-sample t-test, *P<0.05, **P<0.01, ***P<0.001). **(D, E)** MelJuSo expressing cytosolic (NES-GFP)^2^-CL1 (D) and nuclear (NLS-GFP)^2^-CL1 (E) reporter substrates were treated with indicated concentrations GC7 and samples were analyzed by flow cytometry for YFP expression (n=3, mean ± SD, Kruskal-Wallis test, *P<0.05, **P<0.01, ns: non-significant). **(F)** Single clonal EIF5A knockout (EIF5A KO) in MelJuso tBFP-H2A-p2A-YFP-CL1 reporter cells were analyzed by flow cytometry for fluorescence in YFP and tBFP. KO cell lines were plotted for their tBFP versus YFP ratio in comparison to parental reporter cells. **(G)** Two EIF5A KO MelJuSo tBFP-H2A-p2A-YFP-CL1 reporter cells or parental reporter cells with the same tBFP intensities were treated 16 hours with 100 nM epoxomicin and analyzed by flow cytometry for their YFP/tBFP ratio. Data are presented as box plot with median and 5 - 95 percentile. (n=4, paired one-way ANOVA, *P<0.05, **P<0.01).

We next investigated whether the enhanced ubiquitin-dependent proteasomal degradation of YFP-CL1 in eIF5A-deficient cells depends on its cognate ubiquitination pathway. Depletion of the ER-associated ubiquitin conjugase UBE2G2 inhibited the enhanced clearance of YFP-CL1 in GC7-treated cells, indicating an important role for its cognate ubiquitination enzymes (**Fig. 5B, Suppl. Fig. 5A**). The depletion of the two major E3 ligases RNF139 or UBE3C alone did not prevent the effect of GC7 (**Fig. 5C**), which may be due to the observed redundancy of YFP-CL1 targeting ubiquitin ligases, although we cannot exclude the involvement of other ubiquitin ligases. Using cell compartment-specific YFP-CL1 reporters (Bennett et al, 2005), we found that cytosolic localization of the reporter was a prerequisite for enhanced degradation by GC7 (**Fig. 5D**), as the nuclear YFP-CL1 reporter was unresponsive (**Fig. 5E**), consistent with ubiquitination occurring at the ER. Together, these findings suggest that the ubiquitination of YFP-CL1 at the cytosolic face of the ER membrane facilitates the enhanced degradation in eIF5A-deficient cells.

To further validate the role of eIF5A in regulating YFP-CL1 stability, we deleted the EIF5A gene in the YFP-CL1/tBFP-H2A reporter cell line, confirmed by the absence of eIF5A and hypusine reactivity (**Suppl. Fig. 5B**). Although EIF5A knockout clones were viable, they exhibited reduced growth kinetics (**Suppl. Fig. 5C**), as has been reported for EIF5A ablation (Mandal et al, 2016). In line with the expected accelerated degradation of YFP-CL1, we found that independent EIF5A-knockout clones had on average a 30% reduced YFP-CL1/tBFP-H2A ratio compared to the parental cells (**Fig. 5F**). The decreased ratio was consistent throughout all clones tested independently of the expression levels of the reporter based on the fluorescence intensity of the tBFP-H2A reference protein. The enhanced proteasomal degradation of YFP-CL1 in EIF5A-knockout cells was further supported by the observation that treatment with epoxomicin stabilized YFP-CL1 to a larger extent in EIF5A-knockout cells as compared to the parental cells (**Fig. 5G**). We conclude that genetic or chemical inhibition of eIF5A accelerates the proteasomal degradation of YFP-CL1.

### The mitochondrial network regulates YFP-CL1 stability

It has previously been reported that eIF5A inhibition induces a metabolic switch from oxidative phosphorylation to glycolysis, resulting in dramatic changes in the mitochondrial network (Barba-Aliaga & Alepuz, 2022). This, combined with the observation that many mitochondrial proteins were overrepresented among the potential suppressors of YFP-CL1 degradation in our CRISPR/Cas9 screen, suggests that the mitochondrial compartment may regulate YFP-CL1 stability.

To better understand the contribution of the mitochondria in the turnover YFP-CL1, we performed fractionation experiments in which the ER and mitochondrial fractions were separated and probed for the presence of YFP-CL1. YFP-CL1 was readily detected in both ER and mitochondrial fractions, indicating that the amphipathic helical CL1 interacts with both membranous compartments (**Fig. 6A**). Consistent with the ER being the primary site for YFP-CL1 ubiquitination, we observed YFP-CL1 species migrated at higher molecular weights in a ladder-like pattern typical for ubiquitinated proteins in ER fractions of proteasome inhibitor-treated cells. In sharp contrast, only unmodified YFP-CL1 was detected in the mitochondrial fraction even after proteasome inhibition. The importance of eIF5A for the mitochondrial localization of YFP-CL1 was substantiated by the failure to detect the reporter in mitochondrial fractions of EIF5A knockout clones (**Fig. 6B**). This supports a model where the ER and mitochondrial compartments have opposite effects on YFP-CL1 stability, with the ER compartment playing a central role in ubiquitination and the mitochondrial compartment interfering with this process.

**Figure 6:**
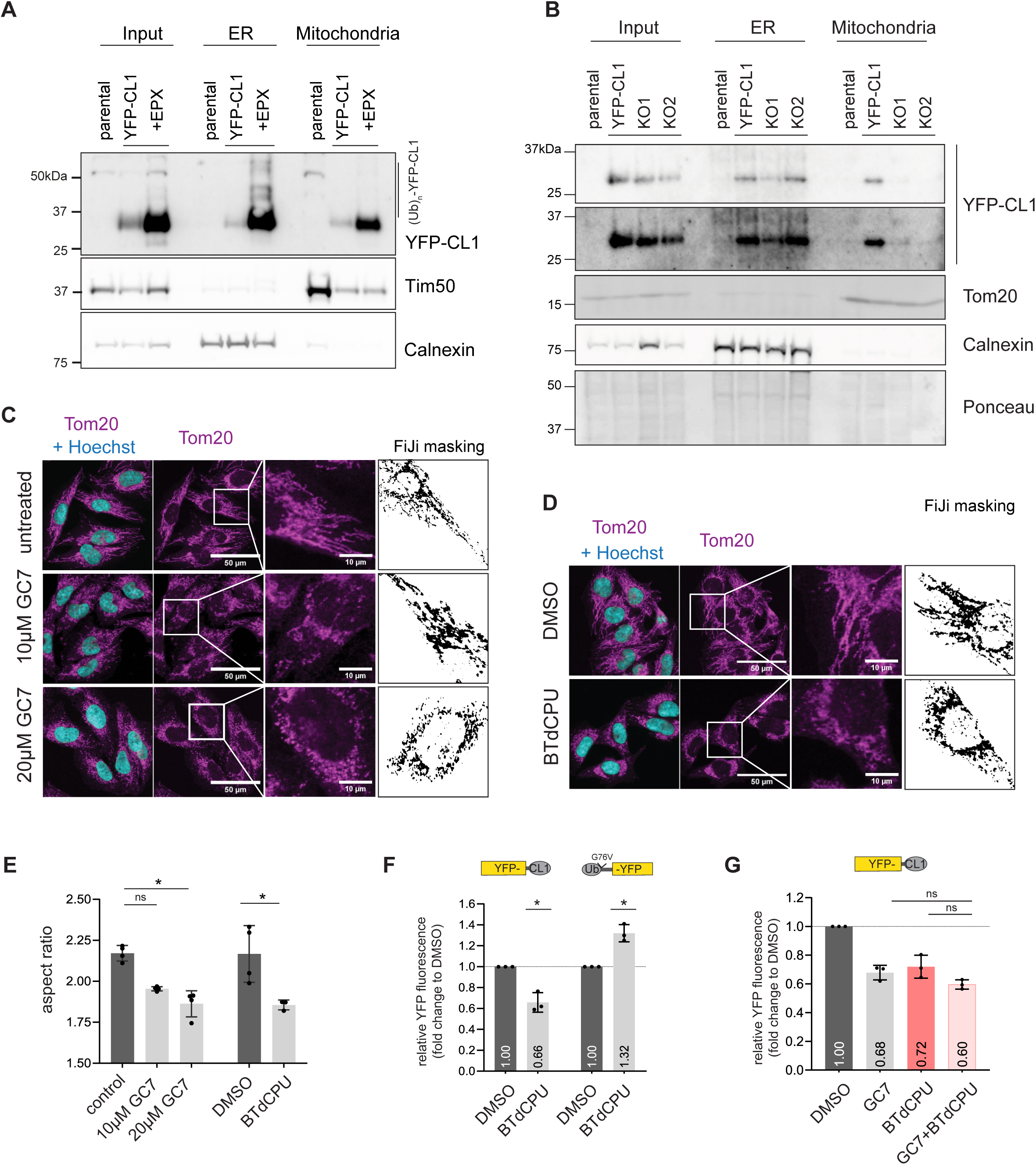
The mitochondrial network regulates YFP-CL1 stability. **(A, B)** MelJuSo dual-fluorescent reporter cells were fractionated and 2.5 µg (A) or 2 µg (B) protein of each fraction sample was analyzed by western blotting using anti-GFP for reporter expression and compartment specific antibodies anti-Tim50 and anti-Calnexin. **(C, D)** MelJuSo cells were treated with 10 µM and 20 µM GC7 (C) or 10 µM BTdCPU (D) for 24 hours, stained for the mitochondrial network with an anti-Tom20 antibody and imaged by confocal microscopy. Representative images, larger image: scale bar = 50 µm, zoom-in image: scale bar = 10 µm. Deconvoluted images, as indicated by a representative cell, were used for staining quantifications. **(E)** Quantification of the length of the mitochondrial filaments per image from (B+C) (n= 4 for GC7 and n=3 for BTdCPU, mean ± SD, Kruskal-Wallis test, *P<0.05, ns: non-significant). **(F)** MelJuSo YFP-CL1 or Ub^G76V^-YFP cells were treated with 10 µM BTdCPU and samples were analyzed by flow cytometry for YFP expression (n=3, mean ± SD, one-sample t-test, *P<0.05). **(G)** MelJuSo YFP-CL1 cells were treated with 10 µM GC7, 10 µM BTdCPU or a combination of both compounds. Samples were analyzed by flow cytometry for YFP expression (n=3, mean ± SD, Kruskal-Wallis test, ns: non-significant).

Consistent with reorganization of the mitochondrial compartment in eIF5A-deficient cells (Melis et al, 2017), microscopic analysis showed that GC7 treatment altered the mitochondrial distribution to a more fragmented network (**Fig. 6C**). Similar changes in the mitochondrial network were also observed in the EIF5A knockout clones compared to the parental cell line (**Suppl. Fig. 6A**), confirming the importance of eIF5A for maintaining an elaborate mitochondrial network. Another compound that affects the integrity of the mitochondrial network is 1-(benzo[d][1,2,3]thiadiazol-6-yl)-3-(3,4-dichlorophenyl)urea (BTdCPU) (Perea et al, 2023). Microscopic analysis confirmed that, like GC7 treatment, BTdCPU administration changed the mitochondrial network connectivity (**Fig. 6D, E**). Importantly, treatment with BTdCPU reduced the levels of YFP-CL1 while having the opposite effect on the Ub^G76V^-YFP reporter (**Fig. 6F**), similar to the effects observed for GC7 treatment (**see Fig. 4C, D**). BTdCPU treatment did not affect eIF5A hypusination (**Suppl. Fig. 6B, C**), consistent with the compound altering the mitochondrial network through an independent mechanism (Perea et al, 2023). Co-treatment of GC7 and BTdCPU had no additional effect on YFP-CL1 stability, indicating that the compounds promote the decrease in YFP-CL1 via a shared mechanism (**Fig. 6G**). This was further supported by the observation that BTdCPU did not further decrease the YFP-CL1 levels in eIF5A-depleted cells **(Suppl. Fig. 6D**). It is noteworthy that the effect of GC7 and BTdCPU on YFP-CL1 degradation is not caused by mitochondrial dysfunction as administration of the mitochondrial complex I inhibitor rotenone did not decrease the steady-state levels of YFP-CL1 (**Suppl. Fig. 6E**). Together, these data support a model in which eIF5A inhibition enhances the degradation of YFP-CL1 by driving mitochondrial network remodeling, ultimately facilitating its release from the mitochondria. Given the pivotal role of the ER compartment in YFP-CL1 ubiquitination and the presence of a pool of mitochondria-associated YFP-CL1 that lacks ubiquitin modifications, this suggests that the dissociation of YFP-CL1 from mitochondria, accompanying the reorganization of the mitochondrial network in eIF5A-deficient cells, increases the degradation-accessible pool of YFP-CL1 and promotes clearance of YFP-CL1.

### Inhibition of eIF5A stimulates degradation of engineered and disease-associated, amphipathic helix-containing proteins

We next investigated whether the enhanced ubiquitin-dependent proteasomal degradation of YFP-CL1 in eIF5A-deficient cells depends on the amphipathic nature of the CL1 degradation signal. Previous studies have shown that amino acid substitutions reducing the hydrophobicity of the amphipathic helix in the CL1 domain decrease ubiquitination by ER-localized ubiquitin ligases (Gilon et al, 2000), and stabilize YFP-CL1 (Stefanovic-Barrett et al, 2018). To explore the importance of the amphipathic nature of the CL1 degradation signal in the enhanced degradation observed in eIF5A-depleted cells, we utilized a CL1 mutant with a distorted amphipathic helix (CL1 ACKN<u>WF</u>SSLSHFVIHL; CL1* = ACKN<u>AA</u>SSLSHFVIHL) (Stefanovic-Barrett et al, 2018) (**Fig. 7A**). As expected, the amino acid substitutions resulted in increased steady-state levels of the YFP-CL1* (**Fig. 7B**) and reduced accumulation of the reporter in response to proteasome inhibition (**Fig. 7C**), reflecting stabilization of the reporter. Microscopy analysis also confirmed that the reduced hydrophobicity was accompanied by a decreased tendency to aggregate into intracellular inclusions upon proteasome inhibition (**Fig. 7D, E**). Importantly, no enhanced degradation of YFP-CL1* was observed in eIF5A-depleted cells (**Fig. 7F**), indicating that the presence of an amphipathic helix in the CL1 degradation signal, which renders the protein aggregation-prone, is also critical for both the constitutive and enhanced degradation of YFP-CL1.

**Figure 7:**
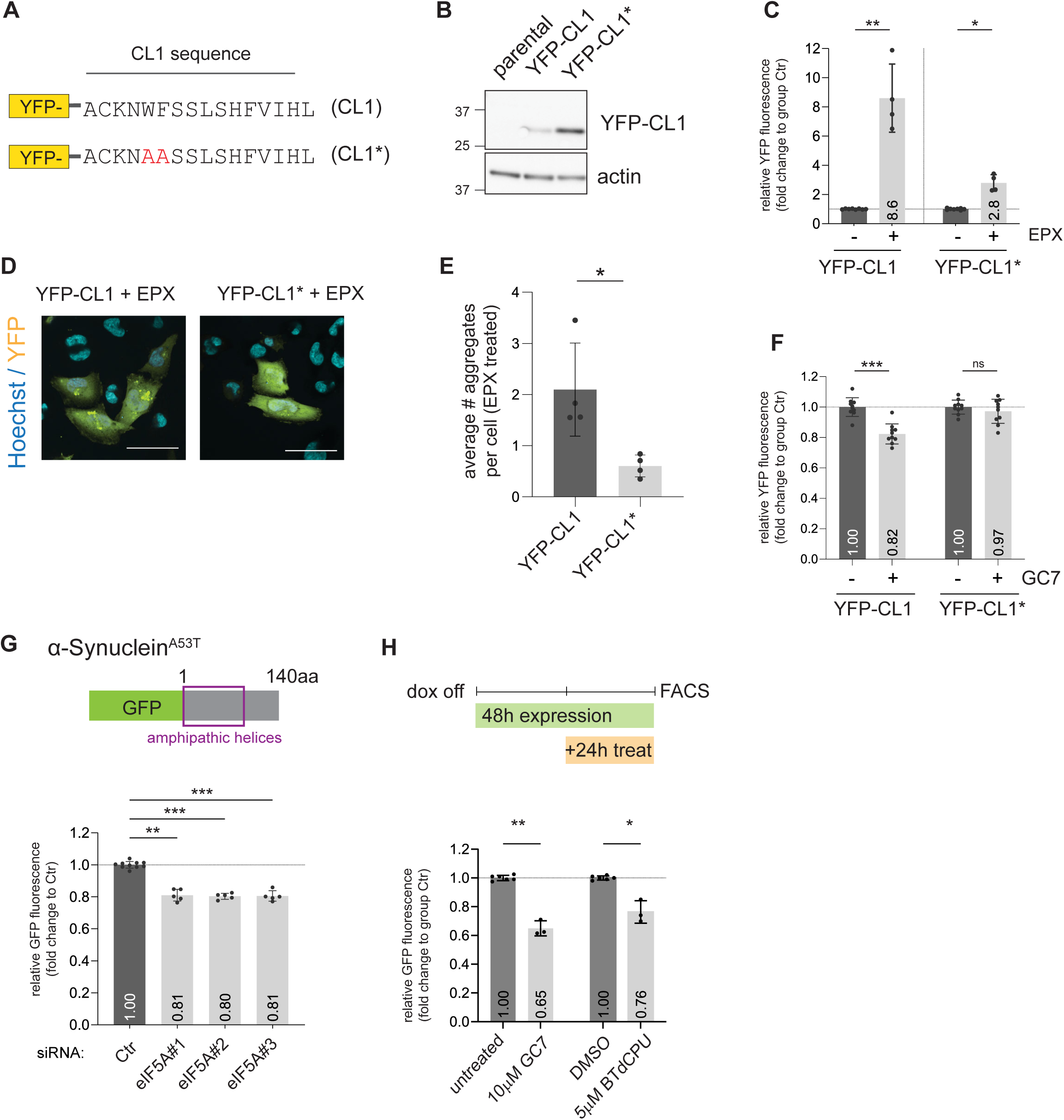
Inhibition of eIF5A stimulates degradation of engineered and disease-associated, amphipathic helix-containing proteins. **(A)** Schematics of the CL1 and CL1* sequence. **(B)** YFP-CL1 and YFP-CL1* were transiently overexpressed in MelJuSo cells for 24 hours and analyzed by western blotting using an anti-GFP antibody. **(C)** YFP-CL1 and YFP-CL1* were transiently overexpressed in MelJuSo cells for 48 hours and treated the last 16 hours with 100 nM epoxomicin (EPX). Samples were analyzed by flow cytometry for YFP expression separately for each mutant. EPX treated samples were set in relation to their respective control (n=4, mean ± SD, Kruskal-Wallis test, *P<0.05, **P<0.01). **(D)** YFP-CL1 and YFP-CL1* were overexpressed in MelJuSo cells for 48 hours and treated the last 16 hours with 100 nM epoxomicin (EPX). Cells were imaged by confocal microscopy. Representative images, scale bar = 50 µm. **(E)** Images were analyzed for the average number of cells with aggregates per condition (n=4, mean ± SD, Mann-Whitney test, *P<0.05). **(F)** YFP-CL1 and YFP-CL1* were transiently overexpressed in MelJuSo cells for 48 hours and treated the last 24 hours with 12 µM GC7. Samples were analyzed by flow cytometry for YFP expression separately for each mutant and set in relation to their respective control (n=5, mean ± SD, Kruskal-Wallis test, ***P<0.001, ns: non-significant). Controls are the same as shown in Fig. 7C. **(G)** MelJuSo cells with inducible expression of GFP-α-synuclein^A53T^ expression were transfected with 20 nM siRNAs for 72 hours and GFP expression was analyzed by flow cytometry (n=5, mean ± SD, Kruskal-Wallis test, **P<0.01, ***P<0.001). **(H)** GFP-α-synuclein^A53T^ expression was induced in MelJuSo cells with inducible for 48 hours, treated the last 24 hours with 10 µM GC7 or 5µM BTdCPU and analyzed by flow cytometry for GFP expression in independent experiments. (n=3, mean ± SD, Kruskal-Wallis test, *P<0.05, **P<0.01).

A prime example of a disease-associated, amphipathic helix-containing protein is α-synuclein, which is linked to Parkinson’s disease and other neurodegenerative diseases (Abou-Sleiman et al, 2006). This aggregation-prone protein contains two amphipathic helices (Trexler & Rhoades, 2009) and localizes to mitochondrial membranes (Reed et al, 2023). To address whether eIF5A regulates also degradation of disease-associated mutant α-synuclein, we generated a stable cell line that allowed inducible expression of GFP-tagged α-synuclein^A53T^. Consistent with the proposed effect of eIF5A deficiency on amphipathic helix-containing proteins, depletion of eIF5A reduced GFP-α-synuclein^A53T^ levels in this stable cell line (**Fig. 7G**). Moreover, administration of GC7 or BTdCPU also resulted in a decrease in the GFP-α-synuclein^A53T^ levels (**Fig. 7H**). Together, these findings show that preventing mitochondrial localization of aggregation-prone proteins can enhance their clearance.

## Discussion

Despite detailed knowledge of aggregation-prone proteins that cause neurodegenerative disease, little can be done to mitigate the devastating consequences for patients suffering from these conditions. As cells are equipped with protein quality control systems that identify, intercept, and destroy aggregation-prone proteins (Sala et al, 2017), leveraging proteasomal degradation using the cell’s innate protective mechanisms may allow selective targeting of disease-associated proteins while restricting collateral damage. The fact that cells under challenging conditions prevent protein aggregation by enhancing the degradation of misfolded proteins as part of stress responses implies that it is possible to stimulate the UPS within the boundaries of its physiological capacity (Flick & Kaiser, 2012).

In agreement with earlier studies (Leto et al, 2019; Stefanovic-Barrett et al, 2018), we found that ER localization and ER protein quality control proteins are the main determinants for efficient proteasomal degradation of the aggregation-prone YFP-CL1 reporter. Consistent with this scenario, sgRNAs targeting genes encoding insertion and removal of proteins from the ER were enriched in the YFP-CL1^high^ and YFP-CL1^low^ populations, respectively, suggesting that increased residence time at the ER enhances degradation of this tail-anchored reporter. Unexpectedly, our CRISPR-Cas9 screen indicated an opposite link between mitochondrial homeostasis and YFP-CL1 degradation, suggesting that mitochondrial localization may hinder efficient clearance. Although the GO analysis showed that sgRNAs targeting genes encoding mitochondrial proteins were most abundant in the sorted population with reduced YFP-CL1 levels, the enrichment was relatively modest, with as striking exception eIF5A, a translation factor that plays a central role in the regulation of mitochondrial homeostasis (Barba-Aliaga & Alepuz, 2022). Further analysis confirmed that genetic or chemical interference with eIF5A function accelerates the degradation of both engineered and disease-associated aggregation-prone proteins.

The tendency of the aggregation-prone CL1 domain to interact with membranous structures, particularly the ER and mitochondria, appears to be critical for the effect of eIF5A on the proteasomal degradation of these proteins. Even though the CL1 motif was originally identified as an ER-binding peptide (Gilon et al, 2000), it also resembles the amphipathic helical motifs that target tail-anchored proteins to other membranous organelles, such as mitochondria (Avendano-Monsalve et al, 2020). Whether the CL1 motif interacts passively with the outer mitochondrial membrane due to its hydrophobic nature or engages the mitochondrial machinery for inserting tail-anchored proteins remains to be determined. Notably, the molecular mechanism responsible for the reduced association of YFP-CL1 with mitochondria in eIF5A-deficient cells remains to be determined. Our data suggest that disturbing mitochondrial homeostasis may not be the sole explanation, given that the mitochondrial complex I inhibitor rotenone did not accelerate the degradation of the reporter. Therefore, specific regulatory mechanisms for amphipathic helix-containing proteins that inhibit the insertion and/or stimulate the extraction of tail-anchored mitochondrial proteins may be involved in the reduced mitochondria association and enhanced degradation in the absence of functional eIF5A. It is noteworthy that a recent study revealed a role for eIF5A in regulating mitochondrial import through facilitating translation of the TIM50 translocase in yeast (Barba-Aliaga et al, 2024).

We propose that the mitochondrial compartment functions as a holdout compartment for aggregation-prone proteins, keeping them out of reach of the primary systems involved in protein quality control, such as those located at the cytosolic face of the ER (**Fig. 8**). Due to the high levels of protein synthesis at the ER in combination with the challenging environment for protein folding, the ER constantly requires an efficient system for dealing with misfolded proteins, as evidenced by the presence of various sensors that mobilize the unfolded protein response (Krshnan et al, 2022). The presence of this efficient system may explain why the cytosolic face of the ER may function at the same time as a local hub for dealing with cytosolic aggregation-prone proteins that tend to associate with ER membrane, due to the intrinsic nature of these domains. Even though localizing the ubiquitination of aggregation-prone proteins at the ER may be beneficial in terms of efficiency, this may also impose limitations, as mislocalization to other membranous organelles may provide an escape route.

**Figure 8:**
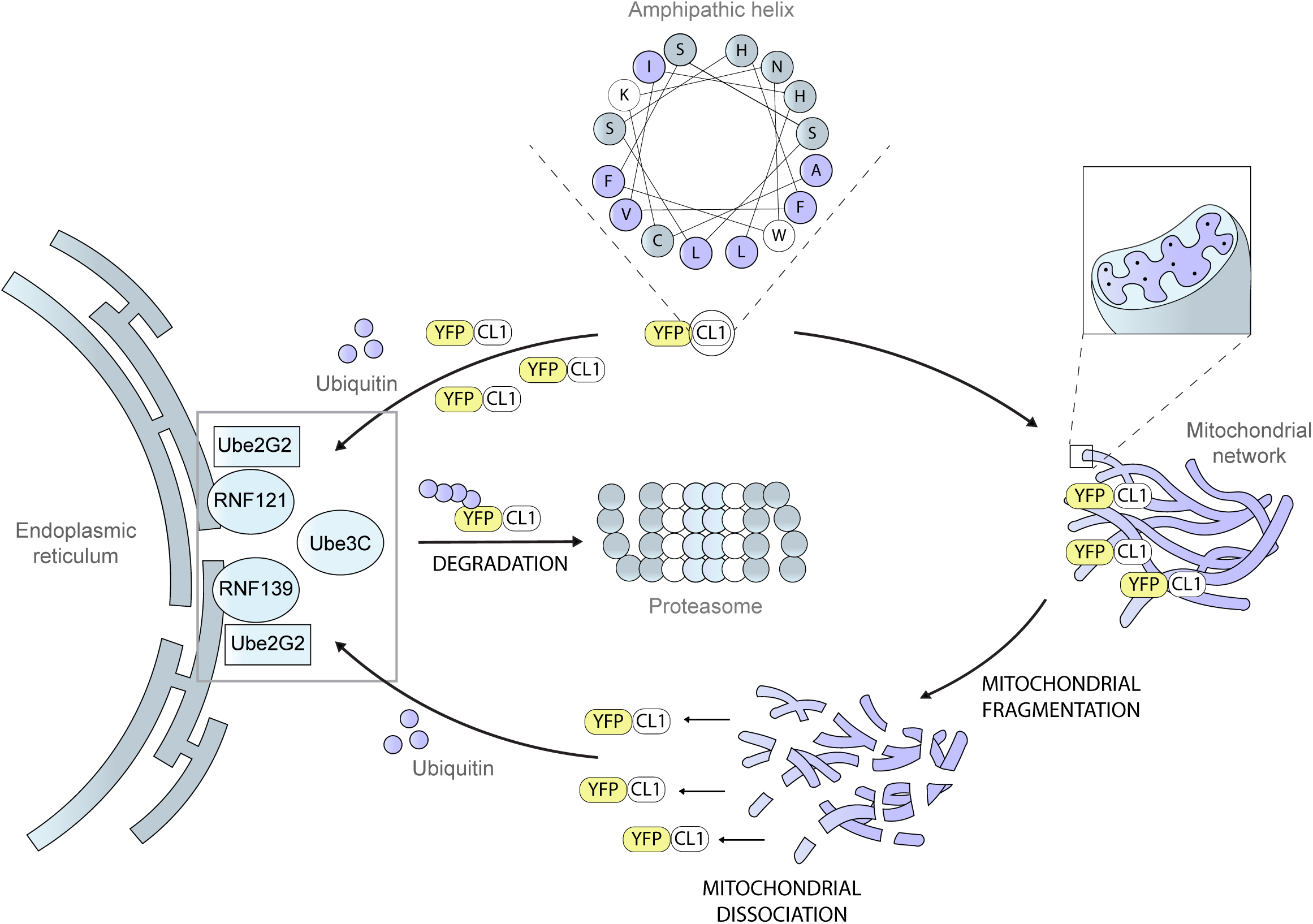
Schematic representation of model. Under steady-state conditions the aggregation-prone YFP-CL1 is either constantly ubiquitinated by several ER-resident E3 ligases and degraded by the proteasome or mistargeted to mitochondria resulting in a long-lived protein pool. Mitochondrial fragmentation caused by eIF5A depletion or inhibition reduces mislocalization of YFP-CL1 and increases the reporter pool that is susceptible to ubiquitinated and degradation.

Recent studies have revealed that eIF5A is also involved in other processes related to protein quality control, suggesting a more integrated role of this translation factor in maintaining protein homeostasis. Notably, eIF5A is critical for CAT tailing (Tesina et al, 2023), an important step in targeting newly synthesized polypeptides for ubiquitin-dependent proteasomal degradation through ribosome quality control (RQC) (Kostova et al, 2017). Additionally, eIF5A-deficient cells exhibit reduced levels of the autophagy protein Atg3 that is required for lipidation of LC3/GABARAP and the formation of autophagosomes, resulting in impaired degradation of proteins through the autophagolysosomal system (Lubas et al, 2018). This suggests that changes in the levels of hypusinated eIF5A can potentially rewire proteolytic systems in response to the metabolic state of cells.

Our finding that eIF5A affects not only synthesis but also degradation of proteins adds another level of complexity to its regulatory role in the cellular proteome. A major challenge in deciphering the effect of eIF5A on the cellular proteome will be to distinguish if reduced steady-state levels of a protein are caused by inhibition of translation, accelerated degradation, or a combination of both. In this context, it is intriguing that even though the reduction in Atg3 levels has been attributed to the presence of a non-canonical tripeptide motif that requires eIF5A for efficient translation (Lubas et al, 2018), it also contains an amphipathic helix important for its interaction with autophagosomes (Nishimura et al, 2023). Therefore, it remains possible that in addition to the effect on translation, the reduced Atg3 levels may also result from accelerated proteasomal degradation in the absence of hypusinated eIF5A.

The question remains whether eIF5A could be a viable therapeutic target to accelerate UPS-mediated clearance of disease-associated proteins in neurodegenerative diseases. The unique dependency of eIF5A on hypusination may provide a promising avenue to chemically fine-tune its activity to accelerate clearance of disease-associated proteins in neurodegenerative diseases. Encouragingly, short-term systemic inhibition of eIF5A hypusination by GC7 has shown protective effects in mouse models for type 2 diabetes (Robbins et al, 2010), renal ischemia (Melis et al, 2017), and stroke (Bourourou et al, 2021). Importantly, inhibition of eIF5A may not be the only means to reach this goal, as alternative ways to interfere with the mitochondrial localization of disease-associated proteins, without affecting mitochondrial homeostasis, may be equally effective in reducing the levels of these proteins and mitigating their toxic effects.

## Material and Methods

### Cell culture and reagents

MelJuSo cells (RRID:CVCL_1403) and all MelJuSo derived stable clonal cell lines were cultured in DMEM with GlutaMAX (Invitrogen) and supplemented with 10% fetal bovine serum (Invitrogen) at 37°C and 5% CO_2_. MelJuSo reporter cells YFP-CL1, Ub^G76V^-YFP, Ub-R-YFP and CD3δ-YFP have been described previously (Menendez-Benito et al, 2005). Cells were routinely tested for Mycoplasma infection. MelJuSo cells have been authenticated by STR profiling 2024/09 (Microsynth). All reagents used in this study can be found in **Suppl. Table 3**. The tet-off stable cell line was cultured in the presence of 0.1 µg/ml doxycycline (Sigma). Where indicated, cells have been treated with the compounds epoxomicin (Sigma), bortezomib (Sigma), GC7 (Sigma), BafA1 (Enzo), cycloheximide (Sigma), TAK-243 (Cayman Chemicals) and BTdCPU (MedChem Express) for the indicated time points in antibiotics-free medium.

### Generation of stable cell lines

Stable cell lines were generated by using selection medium 48 hours post-transfection for 1 week (1.5 mg/ml for G418 (Invitrogen) or 1 μg/ml puromycin (Sigma)). Then, cells were either sorted for positive fluorescent markers as a polyclonal cell line or seeded for single colonies and further grown up, both in the presence of selection medium if needed for an additional two weeks. For the screening cell line, the UPS reporter cells were first generated as a monoclonal cell line and expression was validated by western blotting and flow cytometry. Then, cells were transduced with Cas9 and selected with bleomycin. To generate eIF5A knockout cell lines, the reporter cell line with stable Cas9 expression was transiently transfected with Lipofectamine™ CRISPRMAX™ Cas9 transfection reagent. In brief, the transfection reagent was mixed with the TRUE® synthetic gRNA against EIF5A (CRISPR288707_SGM, Invitrogen) in OptiMEM (Invitrogen) and added to the cells for 48 hours. Then, cells were grown as single colonies and analyzed by western blotting. To generate inducible cell lines, cells were generated as described above. Medium with 0.1 ng/μl doxycycline was changed 6 hours posttransfection and changes every other day. To sort a polyclonal positive cell line, cells were shortly cultured in medium only for 14 hours, sorted and cultured again in the presence of doxycycline.

### CRISPR-Cas9 knockout screen

MelJuSo reporter cells were transduced with the Brunello-UMI library by lentivirus as described previously (Giovannucci et al, 2021). In brief, cells were transduced with a multiplicity of infection of 0.4 and about 1000 cells/guide in 2 μg/ml polybrene. After selection with 2 μg/ml puromycin at days 2–7, cells were further cultured. Two independent transductions were performed and cells were used for downstream fluorescence-activated cell sorting (FACS) at days 10, 11, 16 and 17 (replicate 1) and days 14, 15 and 16 (replicate 2). Cells of the YFP-CL1 fractions and the pools were washed once in PBS and genomic DNA was isolated using the QIAamp DNA Blood Maxi kit (Qiagen). The guide and UMI sequences were amplified by PCR, sequencing was performed on an Illumina HiSeq instrument and the Next Generation Sequencing (NGS) data were analyzed with the MAGeCK algorithm (Li et al, 2014), and by UMI lineage dropout analysis (Schmierer et al, 2017). The gene lists for each replicate can be found in **Suppl. Table 1.**

### Flow cytometry and fluorescence-assisted cell sorting

Cells were harvested, washed in PBS and kept at 4°C until flow cytometry analysis using the BD Canto II or the BD LSR III (both Becton, Dickinson & Company) instruments. Depending on the type of experiment, 20 000-100 000 cells were recorded and data were analyzed by FlowJo v10 (Becton, Dickinson & Company). For sorting, cells were additionally washed once in PBS and filtered through a 0.35 µm cell strainer. The sorters BD Aria III and BD Fusion (both Becton, Dickinson & Company) were used, sorted cells were either collected in PBS or medium and further used for downstream analysis or stable cell line generation.

### Screen analysis and GO enrichment

The two independent replicates were analyzed using the MAGeCK gene lists. Enriched genes were identified with the following criteria: a positive MAGeCK score and a minimum of two gRNAs out of four possible in each data set. Then overlap between the two datasets has been analyzed using Venny 2.1 (Oliveros, J.C. (2007-2015) Venny. An interactive tool for comparing lists with Venn’s diagrams. https://bioinfogp.cnb.csic.es/tools/venny/index.html).

To analyze pathway enrichment, Metascape (Zhou et al, 2019) was used to analyze the gene ontology terms of the selected YFP-CL1^high^ and YFP-CL1^low^ hit lists. The top 16 GO terms were presented as bubble plots over the log of the p-value calculated by Metascape.

### Plasmids and cloning

For side-directed mutagenesis, the QuikChange II kit (Agilent) was used according to manufacturer’s instructions and mutagenesis primers can be found in Supplementary Table 1. NEBuilder assembly (New England Biolabs) was used to generate the double fluorescent screening reporter plasmid by assembling fragments with overhangs generated by PCR of cerulean-histone 2A into the pCMV-YFP-CL1 plasmid digested with Age1 (New England Biolabs). The p2A sequence was produced as an oligo with overhangs for the assembly (Integrated DNA Technologies). From the newly generated plasmid, the pCMV-cerulean sequence was cut out by a double digest of Ase1 and Sca1 (New England Biolabs) and replaced by PCR fragments of pEF1α promoter and tBFP. The digested plasmid and the two fragments were again assembled using NEBuilder and verified by sequencing. The tet-off EGFP-synuclein plasmid was generated by PCR amplifying parts of the backbone of the pCW57.1-MAT2A tet-off plasmid (Addgene #100521), EGFP-synuclein (Addgene #40823) and a puromycin resistance cassette. Assembly of the fragments were carried out using NEBuilder and verified by sequencing. All plasmids and primers used in this study can be found in **Suppl. Table 3.**

### Western blotting

Cells were harvested, washed once in PBS and lysed in either 2x LDS buffer (4x LDS sample buffer, 1x protease inhibitor, 1x NuPage reducing agent, 10 µM MG132, 5 mM NEM in PBS) or RIPA buffer (150 mM NaCl, 50 mM Tris, 1% NP40, 0.1% SDS, 0.5% Na-deoxycholate, 10 µM MG132, 5 mM NEM) and for the latter cleared by centrifugation. Proteins were separated on a 12% or 4-12% BisTris NuPage precast gel (Invitrogen) and transferred to either nitrocellulose or PDVF membranes (both Invitrogen). Membranes were blocked with 5% milk in TBS-Tween 0.1% for 1 hours at room temperature (RT). Primary antibodies were incubated 2 hours at RT or overnight at 4°C followed by 2 hours of 1:10,000 diluted secondary antibody at RT with an IRDye (LI-COR) or HRP-linked (CellSignaling) secondary antibody. A primary antibody list can be found at **Suppl. Table 3**. Protein detection was carried out either by infrared detection using an Odyssey (LI-COR) or ChemiDoc (BioRad) or using ECL (Cytivia) or SuperSignal™ West Femto Maximum Sensitivity Substrate (Invitrogen) with chemiluminescence detection at the ChemiDoc or using X-ray films (Fuji). Uncropped blots are presented in **Suppl. Fig. S7 and S8**.

### Mitochondrial fractionations

Fractionations have been carried out as previously described (Schreiner & Ankarcrona, 2017), and adapted to MelJuSo cells. All buffers can be found in **Suppl. Table 3**. All steps were carried out at 4°C or on ice. In brief, 60×10^6^ cells were collected by trypsinization and washed once in PBS. Cells were pelleted at 800x g for 5 minutes and homogenized in ice-cold BFB1 buffer using an automated Teflon homogenizer. The material was then pelleted twice at 800x g for 5 minutes. The cleared supernatant was subjected to 9,000x g for 10 minutes, the supernatant was used to isolate the endoplasmic reticulum while the pellet contained the crude mitochondrial fraction. To isolate the ER, the supernatant was centrifuged at 20,000x g for 20 minutes and transferred to a fresh tube for an additional centrifugation of 100,000x g for 1 hours. The final ER pellet was lysed in RIPA buffer. To isolate pure mitochondria, the pellet was resuspended gently in BFB2 buffer and centrifuged at 10,000x g for 10 minutes. The pellet was then very gently resuspended in BFB3 buffer and centrifuged again at 10,000x g for 10 minutes. The pellet was then resuspended in BFB4 buffer and layered on a 30% Percoll gradient (Sigma) and subjected to centrifugation at 95,000x g for 30 minutes. The pure mitochondrial fraction was then collected, mixed again with BFB4 buffer and centrifuged at 6,000x g for 10 minutes. The pellet was then lysed in RIPA buffer. For all fractions, protein concentrations were measured by BCA (Invitrogen) and equal µg of protein was loaded for the fractions on one SDS-Page gel and subjected for western blotting. Enrichment of the fractions were validated using compartment specific antibodies.

### SiRNA and plasmid transfections

For transient knockdowns, SilencerSelect siRNAs (Invitrogen) were mixed with RNAimax (Invitrogen) according to manufacturer’s instructions and seeded together with the cells. The medium was exchanged after 24 hours. After 72 hours, cells were analyzed. Knockdown efficiencies have been verified by either western blotting or qRT-PCR. For the siRNA mini-screen, three independent SilencerSelect siRNAs (Invitrogen) per gene were used and only selected hit candidates have been verified for knockdown efficiency. For transient plasmid transfections, lipofectamine 2000 or lipofectamine 3000 (Invitrogen) have been mixed with plasmids and added to previously plated cells. After 24 hours, the medium was changed and cells were analyzed after a total of 48 hours overexpression or placed in selection medium for stable cell line generation. A list of used siRNAs and plasmids can be found in **Suppl. Table 3**.

### Quantitative Real-time-PCR (qRT-PCR)

Total RNA was extracted using RNeasy mini kit (Qiagen) according to manufacturer’s instructions. Complementary DNA synthesis was carried out using the following protocol: total RNA was mixed with Oligo(dT)_18_ primer and dNTPs (both Invitrogen) and incubated 5 minutes at 65°C and 2 minutes on ice. Then, 5x first strand buffer, 0.1 M DTT and RNAseOUT (all Invitrogen) were added and incubated 2 minutes at 37°C. M-MLV-RT was added and incubated for 50 minutes at 37°C. The final cDNA was diluted to 10ng/µl and used for qRT-PCR. Quantitative RT-PCR was performed using TaqMan gene expression master mix (Invitrogen) and assays on demand (Applied Biosystems) with the following primers: Hs00183680_m1 (Rnf139), HS02786624_g1 (Gapdh), and Hs01553223_m1 (Rnf121). Cycle threshold (Ct) values were normalized to GAPDH and relative expression values were calculated using the ΔΔCt method.

### Immunofluorescence and imaging

MelJuSo cell lines were grown overnight on coverslips and treated as indicated. Cells were fixed using 4% paraformaldehyde (Invitrogen) for 10 minutes, permeabilized using 0.2% Triton-X100 in 1x PBS for 20 minutes, 100 nM glycine was added for 10 minutes, followed by 3% BSA (Sigma) in 1x PBS for 30 min. The primary antibody, anti-Tom20, was diluted in 0.1% Tween-20 in PBS and incubated overnight at 4°C. It followed a 2-hour incubation at RT with goat anti-rabbit IgG coupled to AlexaFluor 647 (Invitrogen) in 0.1% Tween-20 in PBS. Nuclear staining was performed using Hoechst 33342 (Molecular Probes) 1:5000 in PBS for 30 min. Fixed cells were examined with a Nikon confocal spinning disk microscope (60x/1.42 oil objective). Image processing was performed with FiJi and quantitative analyses were performed using CellProfiler software. For dual-fluorescent reporter cells, MelJuSo cells were grown overnight on coverslips and treated as indicated. Cells were fixed using 4% paraformaldehyde (Invitrogen) for 10 min and washed three times before mounting. Fixed cells were examined with a Leica Stellaris 5 confocal laser scanning microscope (63x/0.75 air objective). Image processing was performed with FiJi.

### Time-lapse imaging and growth analysis

MelJuSo cell lines were seeded in a 6-well plate (Sarstedt) and imaged with the IncuCyte S3 (Sartorius) with constant 37°C and 5% CO_2_. Images were taken at time intervals of 2 hours, for 24 hours. For each well, 4 different sites were imaged. Cell confluency was quantified with the IncuCyte software (2019B Rev2 version).

### Aggregates imaging and quantification

Cells were transfected with the indicated plasmids, after 6 hours detached from the plate and re-seeded on coverslips. Forty-eight hours after transfection, cells were fixed using 4% paraformaldehyde (Invitrogen) for 10 minutes, and nuclei were stained using Hoechst 33342 (Molecular Probes) 1:5000 in PBS for 30 minutes. Fixed cells were examined with a Leica Stellaris 5 confocal laser scanning microscope (20x/0.75 air objective). Four images were taken and cells with aggregates were counted per expression construct and per experiment and related to the total number of nuclei per image.

### Statistics

Statistical analyses were performed using Prism GraphPad v10. Tests are indicated for each experiment in the figure legend; mostly Kruskal-Wallis test with Dunn’s multiple comparison test or one-sample t-test unless otherwise stated. Error bars present +/- SD (standard deviation) unless otherwise stated. The following standard deviations were considered significant: *P≤0.05, ** P≤0.01, ***P≤0.001, **** P≤0.0001.

### AI-assisted technology

Copilot was used to check for spelling and grammatical errors and text editing. The authors take full responsibility for the content of the manuscript.

## Acknowledgements

We thank Maria Masucci and the members of the Dantuma lab for helpful input and the Biomedicum Imaging Center, the CRISPR Functional Genomics Facility and the Biomedicum Flow Cytometry Core Facility for their assistance. We thank Lukas Mann for help with cloning the tet-off plasmid. This work was supported by the Swedish Research Council (N.P.D. 2021-02562), the Swedish Cancer Society (N.P.D. CAN 211653Pj), the Swedish Brain Foundation (FO2022-0271, F2023-0376) and the Karolinska Institute (KID grant). M.E.G. was supported by research fellowships from the Deutsche Forschungsgemeinschaft (DFG) (GI-1329/1-1). N.P.D. is a member of the COST network ProteoCure.

## Author contributions

**M.E.G., N.P.D.** conceptualized the study; **M.E.G, E.B., M.M., M.H.W.** performed the experiments; **L.N.** assisted in the fractionation experiment; **F.A.S.** assisted in microscopy and image analysis; **M.E.G, E.B., M.M., M.H.W., M.A., L.N., F.A.S., N.P.D.** analyzed the data; **M.E.G., N.P.D.** wrote the manuscript draft; **M.E.G., N.P.D.** coordinated the project; All authors edited and approved the final manuscript.

## Disclosure and competing interests statement

The authors declare that they have no conflict of interest.

## Data Availability

This study does not contain deposited data. Original data can be provided upon request.

## Notes

### Competing Interest Statement

The authors have declared no competing interest.

