## Supplementary Figures for "Mitochondria serve as a Holdout Compartment for Aggregation-Prone Proteins hindering Efficient Ubiquitin-Dependent Degradation"

**Supplementary Figure 1: (A,B)** Stable dual-fluorescent reporter cells were plated on cover slips for 48 hours and treated the last 16 hours with 100 nM epoxomicin (EPX) (A) or the last 6 hours with 10 μM TAK-243 (B) and imaged by confocal microscopy. Representative images, scale bar =10μm.

**Supplementary Figure 2: (A)** MelJuSo YFP-CL1 reporter cells were transiently transfected with an siRNA targeting UBE2G2 for 72 hours and analyzed by western blotting for knockdown efficiency. **(B)** MelJuSo YFP-CL1 reporter cells were transiently transfected with an siRNAs targeting RNF139 for 72 hours and analyzed by qRT-PCR for knockdown efficiency. **(C, D)** MelJuSo YFP-CL1 reporter cells were transiently transfected with an siRNAs targeting UBE3C or RNF126 for 72 hours and analyzed by western blotting for knockdown efficiency. **(E)** MelJuSo cells were stained with an anti-RNF121 antibody and imaged by confocal microscopy. Representative image: scale bar = 50 μm, zoom-in image: scale bar = 15 μm. **(F)** MelJuSo cells were fractionated and 10 μg protein of input and ER fraction were analyzed by western blotting using an anti-RNF121 antibody and compartment specific antibodies anti-Tim50, anti-Calnexin and anti-histone H3. Relates to flow cytometry data from Fig. 2D. **(G, H)** MelJuSo YFP-CL1 reporter cells were transiently transfected with siRNAs targeting RNF121 for 72 hours and analyzed by western blotting (G) and qRT-PCR (H) for knockdown efficiencies. **(I, J)** MelJuSo YFP-CL1 reporter cells were transiently transfected with a combination of one or two siRNAs for 72 hours and analyzed by western blotting for knockdown efficiencies with the indicated antibodies.

**Supplementary Figure 3: (A-C)** MelJuSo YFP-CL1 reporter cells were transiently transfected with siRNAs for 48 hours and analyzed by flow cytometry for YFP expression. (n=3 (A, B), mean ± SD, one-sample t-test, *$P<0.05$, ns: non-significant; n=2 (C)). Samples were also analyzed by western blotting for knockdown efficiencies of the siRNAs with the indicated antibodies.

**Supplementary Figure 4: (A)** MelJuSo YFP-CL1 reporter cells were transiently transfected with siRNAs for 72 hours and analyzed by western blotting for knockdown efficiencies with the indicated antibodies. **(B)** MelJuSo YFP-CL1 reporter cells were treated for 24 hours with increasing concentrations of GC7 and analyzed by western blotting for hypusine depletion using indicated antibodies. **(C)** MelJuSo YFP-CL1 reporter cells were transiently transfected with siRNAs for 72 hours and treated the last 24 hours with two different concentrations of GC7. Samples were analyzed by western blotting for knockdown efficiencies and hypusine depletion with the indicated antibodies.

**Supplementary Figure 5:**

**(A)** MelJuSo YFP-CL1 reporter cells were transiently transfected with an siRNA directed against UBE2G2 for 72 hours, treated the last 24 hours with 12 µM GC7, and analyzed by western blotting for knockdown efficiencies and hypusine depletion with the indicated antibodies. **(B)** EIF5A KO cells were established in a dual-fluorescent reporter cell line with stable Cas9 expression using an gRNA at the indicated position. Several clonal cell lines were analyzed for eIF5A depletion by western blotting. **(C)** Two eIF5A KO cell lines and MelJuSo control cells were monitored for their growth curves for 24 hours using confluency measurement (4 images per condition, mean ± SD.

**Supplementary Figure 6:**

**(A)** MelJuSo eIF5A KO and parental cells were stained for the mitochondrial network with an anti-Tom20 antibody and imaged by confocal microscopy. Representative images: scale bar = 100 µm, zoom-in images: scale bar = 25 µm. **(B)** MelJuSo YFP-CL1 cells were treated with 10 µM GC7, 5 µM BTdCPU or a combination of both compounds. Samples were analyzed by western blotting for hypusine depletion with the indicated antibodies. **(C)** Quantification of hypusine levels from western blot data (n=2). **(D)** MelJuSo YFP-CL1 cells were transfected with 20 nM siRNAs against eIF5A for 72 hours, treated the last 24 hours with the 5 µM or 10 µM BTdCPU and analyzed by flow cytometry for YFP fluorescence. (n=3, mean ± SD, one sample t-test when compared to Ctr or Kruskal-Wallis test when samples were compared within each siEIF5A condition, *P<0.05, **P<0.001, ***P<0.0001, ns: non-significant). **(E)** MelJuSo YFP-CL1 cells were treated with 200 nM or 400 nM rotenone for the indicated time points. Samples were analyzed by flow cytometry and one representative histogram is shown of at least two independent experiments.

Supplementary Fig. 1
Gierisch et al.

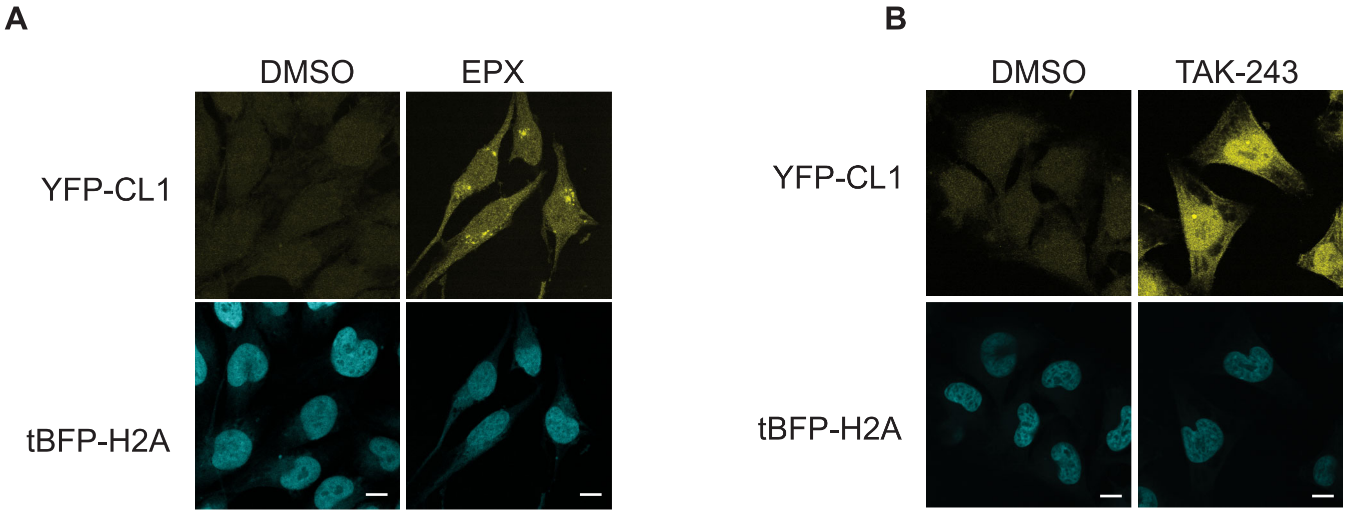

Supplementary Fig. 2
Gierisch et al.

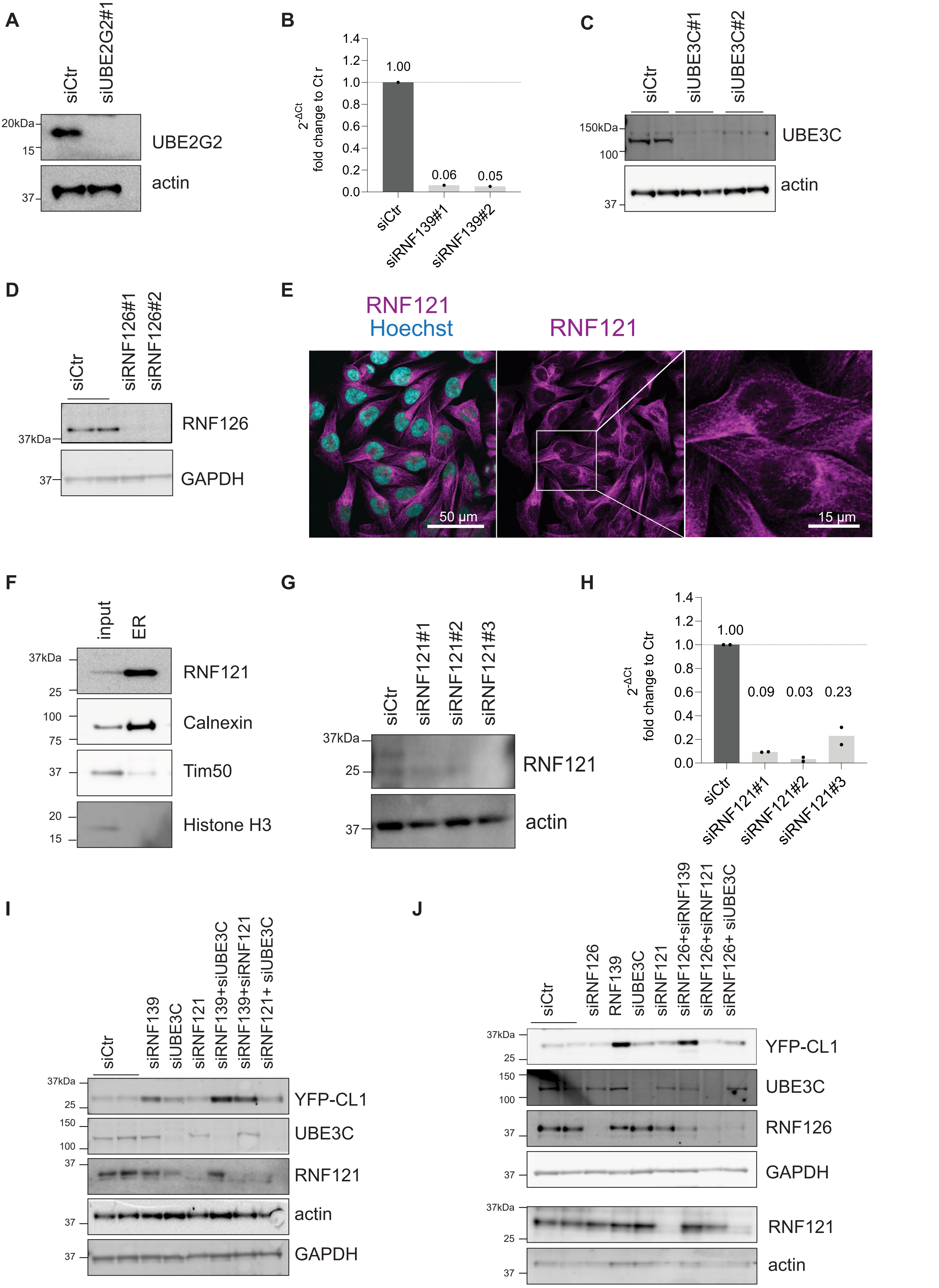

Supplementary Figure 3
Gierisch et al

A

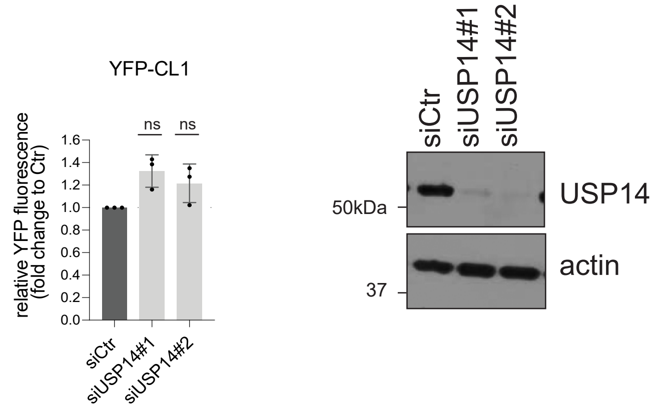


B

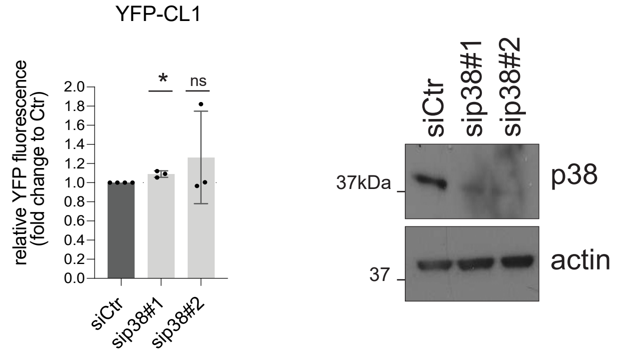


C

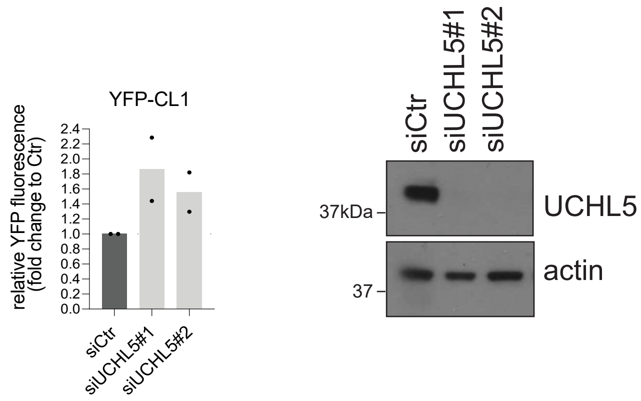

Supplementary Figure 4
Gierisch et al.

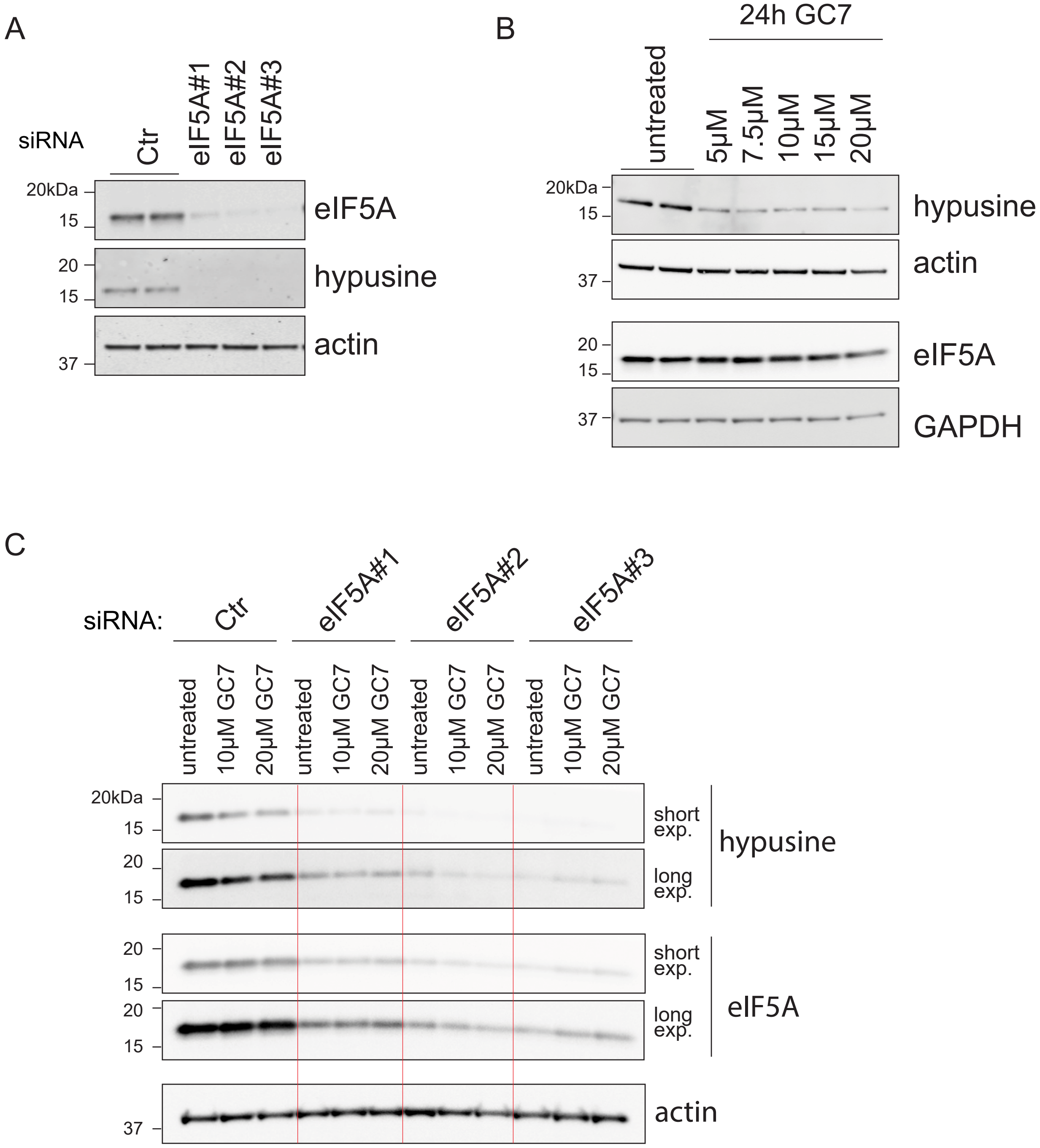

Supplementary Figure 5
Gierisch et al.

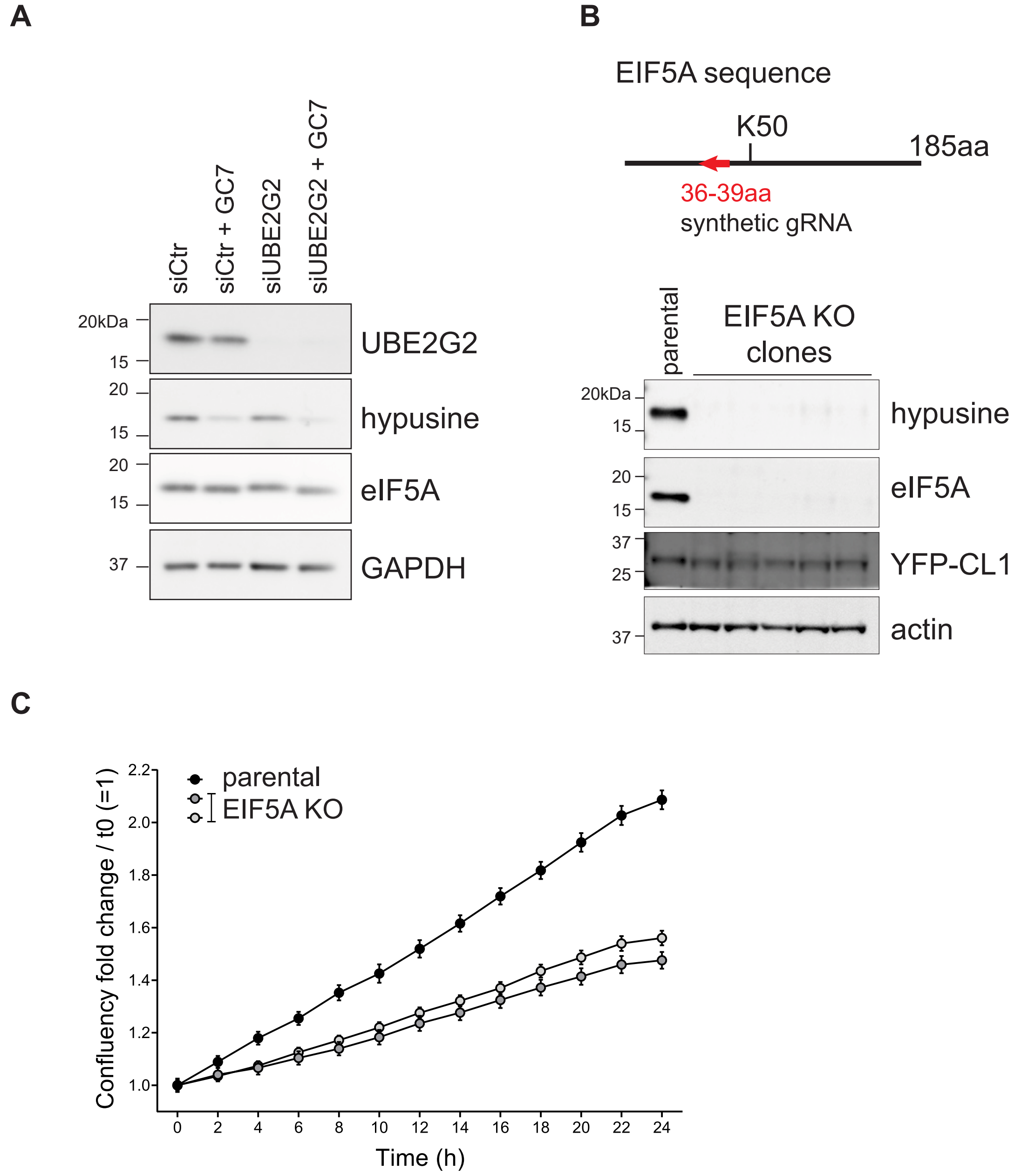

Supplementary Figure 6
Gierisch et al.

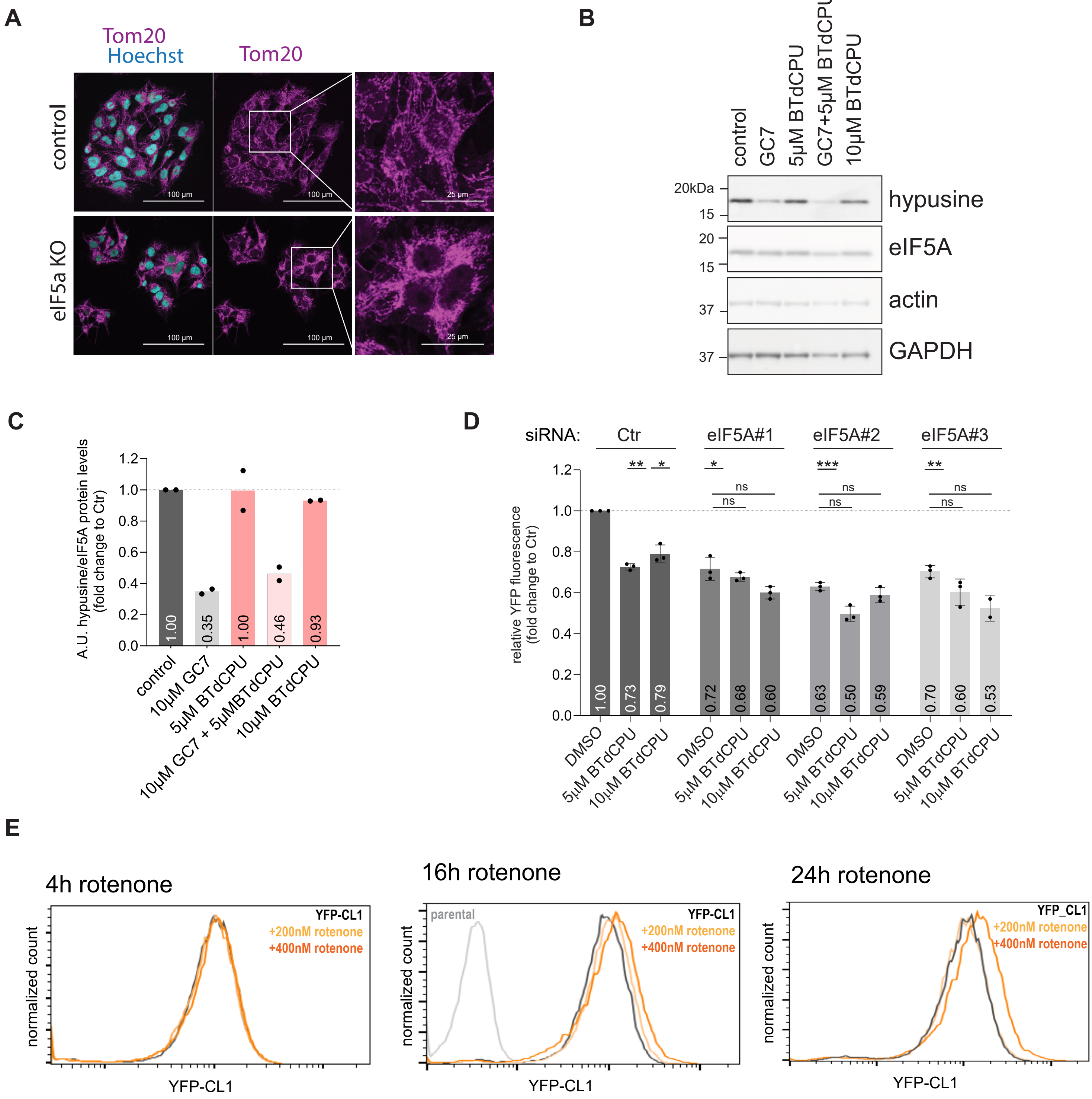
